# Protein Deimmunization with a Generative Diffusion Model

**DOI:** 10.64898/2026.10.09.758136

**Authors:** Zijie Chen, Chih-Chieh (Jay) Yu, Cai Zhou, Xinyun Jiang, Long T. Nguyen, Cameron Myhrvold, Tommi Jaakkola, Kaiyi Jiang

**Affiliations:** Omenn-Darling Bioengineering Institute Princeton University Princeton, NJ, 08544, United States of America; Department of Molecular Biology Princeton University Princeton, NJ, 08544, United States of America; Department of Electrical Engineering and Computer Science Massachusetts Institute of Technology Cambridge, MA, 02139, United States of America; Department of Chemical and Biological Engineering Princeton University Princeton, NJ, 08544, United States of America; Department of Chemistry Princeton University Princeton, NJ, 08544, United States of America; Princeton Language and Intelligence Princeton University Princeton, NJ, 08544, United States of America

**Author notes:** These authors contributed equally (Zijie Chen, Chih-Chieh (Jay) Yu). **Correspondence to**: Kaiyi Jiang.

## Abstract

Protein immunogenicity is a major cause of biologic drug failure, leading to the production of anti-drug antibodies (ADA) that accelerate drug clearance, rejection of repeated doses, abolished efficacy, and sometimes life threatening anaphylaxis in patients. These responses constrain the clinical use of synthetic protein therapeutics from non-human origin. To efficiently deimmunize protein therapeutics, we developed a generative discrete diffusion model (MESSI), which takes protein structures as input and outputs sequences that preserve structure without containing peptides predicted to bind major histocompatibility complex class II molecules. MESSI designs sequences with high structural recovery and down to 0 predicted epitopes for specified MHC molecules at higher efficiency than post-hoc filtering approaches. Experimentally, we first applied MESSI to deimmunize a luciferase and a de novo designed mini-binder against PD-1, resulting in variants with no predicted epitopes and higher activity than their parents. Lastly, we coupled MESSI with evolution guidance to deimmunize uricase, a clinically approved enzyme with severe immunogenicity, and found deimmunized variants with higher catalytic activity than their natural orthologs. MESSI enables generative design of functional proteins with minimized immunogenicity, potentially making protein therapeutics safer and more effective.

## Introduction

Protein therapeutics including antibodies and enzymes are cornerstones of modern medicine (*1*, *2*). In contrast to small molecule based drugs, one of their central liabilities is the immunogenicity of the protein (*3*). Therapeutic proteins are internalized by human antigen presenting cells (APC), processed into peptides, and presented on major histocompatibility complex (MHC) class II molecules on the cell surface of APCs (*4*). Any “non-self” peptide presented could trigger a cascade of reactions leading to the production of anti-drug antibodies (ADAs). ADAs can accelerate clearance, neutralize activity, abolish possibility of repeat dosing, and in the worst cases provoke infusion reactions and life-threatening anaphylaxis (*5*, *6*). The clinical consequences are widespread with documented adverse effects in drugs including interferon beta therapy in multiple sclerosis (*7*, *8*), factor VIII in hemophilia A (*9*), bacterial asparaginase in pediatric leukemia (*10*, *11*), and recombinant uricase in chronic gout patients (*12*, *13*).

The dominant route to a potent and durable ADA response is T cell-dependent (*3*). Therapeutic protein is processed into short peptides in APCs and edited onto MHC class II molecules by human leukocyte antigen (HLA) DM for presentation (*14*). Surface displayed peptide–MHC complexes are surveyed by CD4^+^ T cells, and immunogenic epitopes on the protein could activate the CD4^+^ T cell whose help licenses class-switched, affinity-matured B cell responses (Fig. 1) (*15*, *16*). The crystal structures of HLA-DR established that the class II groove is open at both ends and grips a nine-residue core with flanking extensions, so the actual displayed peptide length is unconstrained from 12 to 25 amino acids and the binding register is not fixed by the peptide termini, in contrast to short and more defined class I peptide (*17*, *18*). To add to this complexity, the MHC class II region is highly polymorphic, so to fully deimmunize a protein, peptides binding to diverse class II molecules have to be considered (*19*–*22*).

**Figure 1.**
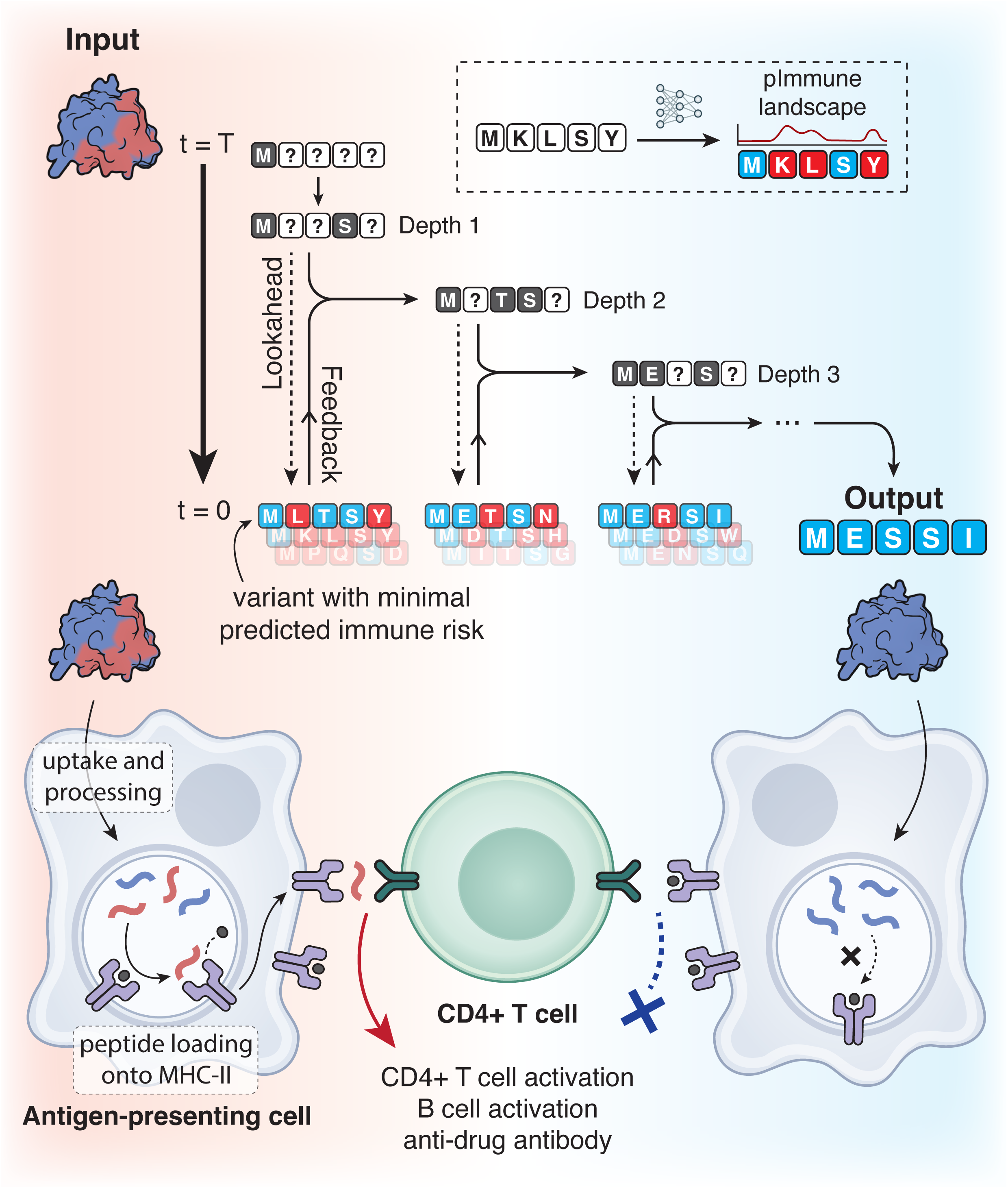
Schematic of MESSI. The top plot shows the model architecture and workflow of MESSI’s guided discrete diffusion model. MESSI takes the 3D structure of a protein as input and fills in amino acid sequences via stepwise diffusion. During each iteration, the model performs a lookahead operation to project a partial sequence into multiple completed sequences, and then estimates the whole-protein-level and local-residue-level immune risk (T cell epitope burden; denoted as pImmune landscape) with a trained immune risk predictor for a particular MHC class II allele (dashed box on upper right). Next, during the feedback operation, the trajectory with the lowest protein-wide immune risk is selected to guide the evolution of the partial sequence, leading to potential adoption of a low-immune-risk amino acid for an open position, or re-opening of a previously determined position due to high immune-risk resulting from the current iteration. The bottom plot illustrates a core pathway leading to anti-drug antibodies against an injected therapeutic protein. Following drug injection, antigen-presenting cells endocytose therapeutic proteins and process them into peptide fragments in endosomes. Peptides with high affinity to MHC-II can displace the invariant starting peptide inside the MHC-II binding groove (shown in dark gray), leading to cell-surface display of the foreign peptide and subsequent anti-drug immune responses. MESSI depletes peptide sequences that are predicted to bind to MHC-II, which can potentially lead to attenuation or inactivation of this anti-drug pathway.

Due to the complexity of the adaptive immune system, even highly humanized antibody drugs can break tolerance and trigger ADA due to the presence of T cell epitopes (*23*). For non-human origin enzymes and *de novo* computationally designed proteins, they typically sit at the far end of the immunogenicity risk distribution due to the fact that their constituent peptides have never been tolerated. Given the rise of generative protein design models recently (*24*–*30*), we are now in an unprecedented time to design novel protein drugs against proteins of interest with speed and precision. However, to realize the potential for these *de novo* designed drugs, their immunogenicity risk has to be addressed upfront in the design stage to make an impact in the clinic.

Previous computational deimmunization tools mostly work by first identifying predicted MHC-II binders in the target protein sequence (*31*–*33*) and then generating mutations to filter for those that break the binding while predicted to preserve global protein structure (*34*, *35*). Episweep and other integer-programming formulations based on Rosetta have been proposed to reduce T cell epitopes while retaining stability or structure (*36*–*40*). These models generally have two key constraints that limit their utility and efficiency when generalizing to diverse proteins and generative design pipelines. First, protein structure and function are often predicted indirectly through structural energies or evolutionary sequence statistics which provide incomplete estimates of the effects of sequence changes especially for large edits. Favorable computational scores therefore often fail to correlate with function, rendering extensive experimental screening and additional optimization necessary (*34*, *39*, *40*). Second, the applicability of individual methods depends on the availability of suitable target-specific information, rendering many methods ineffective for proteins without diverse homologous sequences or prior knowledge to define permissible sites of change. This also makes these models hard to integrate to the modern generative protein design pipeline (*35*, *37*). For methods based on exact combinatorial optimization, an additional limitation is the need to formulate scoring functions and sequence choices in a computationally tractable form. Although these methods can identify Pareto-optimal solutions within their specified models, incorporating more complex predictors or expanding the search space can make exact optimization substantially more demanding (*35*, *40*). In light of these constraints, the field still lacks an efficient and robust model that can be easily integrated into the modern generative protein design pipeline to perform deimmunization for diverse proteins.

For protein structure and function prediction, deep learning-based structural prediction models and generative models have transformed the protein design space (*28*, *41*–*44*). In addition to forward structural prediction models, structure-conditioned inverse folding models, including ProteinMPNN and ESM-IF, are developed to sample from the distribution of sequences compatible with a given backbone, thus rendering them routine tools for enzyme and binder engineering (*27*, *45*–*51*). In particular, language model informed inverse folding approaches, including those based on versatile protein language models such as diffusion protein language models (DPLM) and ESM-IF have shown success in protein engineering in diverse settings, likely due to the incorporation of evolutionary scale sequence information during pretraining (*34*, *45*–*49*, *51*–*53*). The inherent ability to sample diverse sequences for a specified structure with retained or improved function thus presents an unprecedented opportunity to leverage diffusion language model based inverse folding to perform constrained design of potentially non-immunogenic sequences that are depleted of peptides predicted to bind to MHC class II molecules.

Inverse folding models are generally trained to maximize sequence and structure recovery, so controlling and guiding the generation is the critical component in deimmunization since the objective does not align with pretraining tasks. Prior work has attempted to steer the decoding of an autoregressive model away from MHC class I visibility, but only with moderate success and lack of experimental validation (*54*, *55*). Encouragingly, there has been tremendous progress in inference and test time scaling, steering methods for generative models, and discrete diffusion models lately (*56*–*73*). In light of these advances, we developed MESSI: a new generative model to dei**M**munize prot**E**in with di**S**crete diffu**SI**on by steering a frozen structure-conditioned discrete diffusion generator with complete-sequence feedback at inference-time (Fig. 1). MESSI takes a protein structure as input and designs sequences that are predicted to preserve structure while depleting peptides predicted to bind a specified MHC class II molecule (T cell epitopes). MESSI evaluates predicted immune risk (T cell epitopes burden) on complete discrete sequences and uses selected immune risk information together with the corresponding source state to revise the ongoing generation process. MESSI produces designs of high structural confidence carrying as few as zero predicted epitopes for specified MHC-II alleles. Although predicted T cell epitope burden is not the sole factor contributing to clinical immunogenicity in humans because protein aggregation propensity and stability, patient’s immune system status, and the presence of B cell epitopes on the protein also play a role, previous studies have shown a strong correlation of predicted T cell epitope burden with actual ADA incidence and clinically observed immunogenicity (*31*, *32*) and the ability to reduce immunogenicity through deleting predicted T cell epitope (*40*, *74*). In light of this, here we focus on addressing this dominant immune pathway to a durable ADA response and present extensive computational and experimental validation to support MESSI’s use in generative deimmunization.

## Results

### Immune risk feedback guided discrete diffusion

A core event in the development of anti-drug responses is the binding of peptides from the therapeutic protein to MHC-II molecules. Removal of MHC-II epitopes from the protein’s amino acid sequence can reduce immunogenicity, as established by previous studies (*31*–*34*, *40*, *74*). However, balancing the design objective between maintaining protein structure and function and deleting epitopes is a non-trivial task for generative models that excel protein structural prediction. This is partially because immune risk information is needed during the continuous sequence generation process in order to steer its trajectory. However, accurate estimation of immune risk can only be performed on completed discrete amino acid sequences, which is obtained at the end of the diffusion trajectory. Given this challenge, one strategy is to simply select the completed sequences with the lowest estimated immune risk from a large number of sequences generated without any guidance, but this strategy is inefficient and suboptimal in the level of deimmunization.

A better solution is to incorporate the immune risk information into the generation process: retaining favorable amino-acid choices and redesigning positions associated with increased risk. Such feedback would allow each round of design to build on the sequence information identified in earlier rounds. In light of this, we developed MESSI to feedback the immune risk information to an editable, structure-conditioned discrete diffusion process. This feedback first requires rapid, allele-specific evaluation of both whole-protein-level immune risk burden and its local-residue-level risk distribution. We therefore first trained an immune risk predictor on MHC-II eluted-ligand data to provide a continuous, residue-level score for candidate selection and a local risk landscape for sequence revision. Immune Epitope Database (IEDB) contains 364,025 HLA class II eluted ligand records, spanning 125 high-resolution alleles or allele-pair annotations (*75*). For the highly diversified HLA-DRB1 protein, we found 140,010 records with unambiguous, high-resolution single-allele DRB1 annotations (Fig. 2A). For proof of concept, we first focused on three well-represented alleles (HLA-DRB1*04:01, 07:01, and 15:01) that cover a significant portion (∼25%) of global HLA-DR diversity and have abundant mass spectrometry eluted ligand data (*75*). We reasoned that eluted ligand data contain implicit information about how a protein is processed into peptides during proteolysis in APCs in addition to binding affinity to the target HLA-DR proteins, making it an ideal dataset for training (*33*). This will in turn maximize our estimator’s ability to accurately predict peptides that could be processed and presented on class II molecules. The predictor uses a lightweight multiscale dilated convolutional network to encode the protein sequence and score overlapping peptide windows of 12–25 residues in their sequence context. Each allele-specific model is trained jointly with peptide-window supervision and residue-level supervision derived from annotated eluted-ligand coverage. Scores are aggregated over windows covering each residue to construct a local risk landscape, and over all windows to obtain a whole-protein risk score. These complementary summaries support local sequence revision and complete-candidate selection, respectively (Supplementary Text).

**Figure 2.**
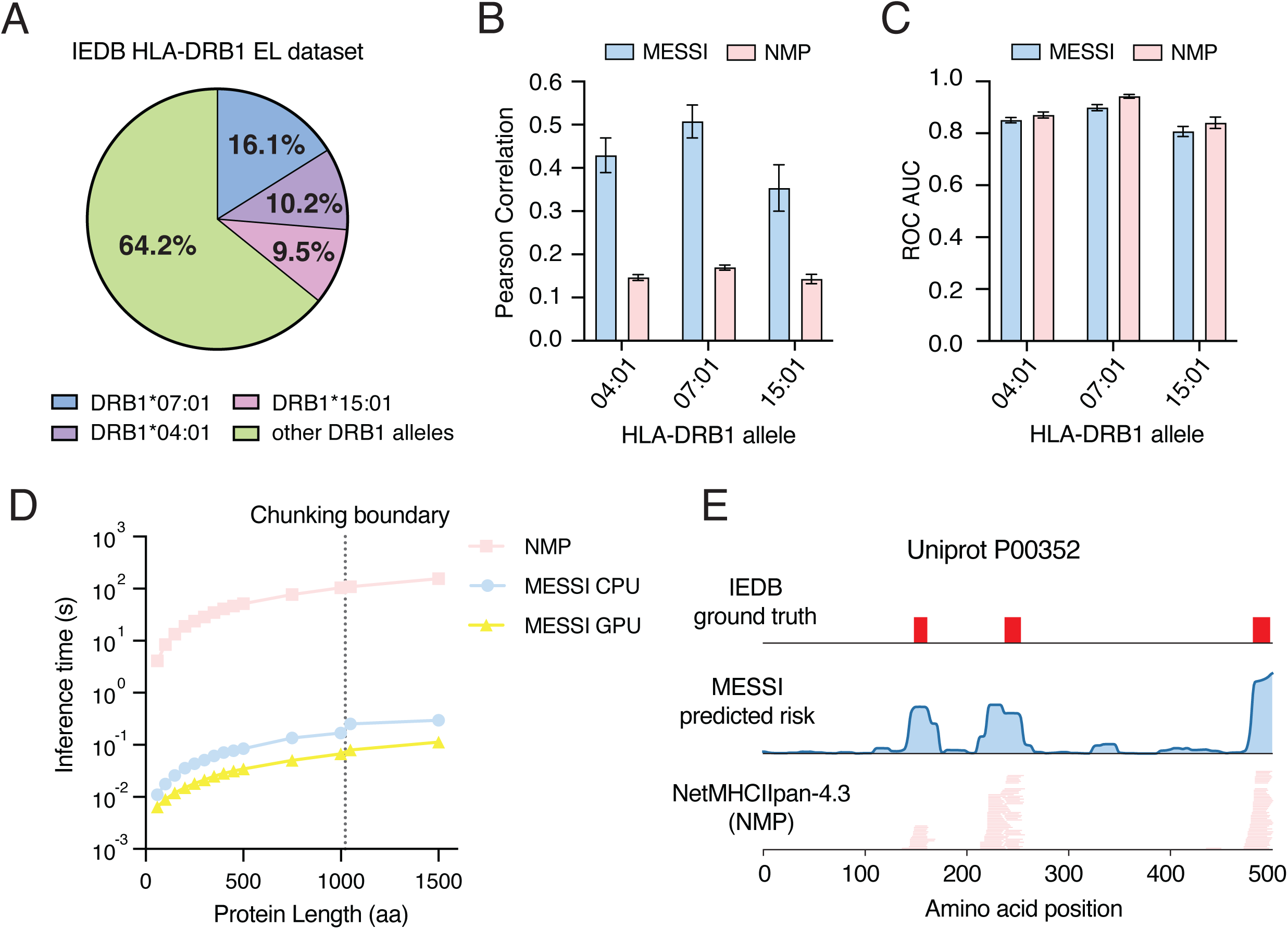
Continuous residue level immune risk estimator. **(A)** Allele composition of the curated IEDB-derived HLA-DRB1 eluted-ligand dataset. The analysis includes 140,010 records with unambiguous, high-resolution single-allele DRB1 annotations, covering 28 alleles. HLA-DRB1*07:01, HLA-DRB1*04:01 and HLA-DRB1*15:01 account for 16.1%, 10.2% and 9.5% of these records, respectively. Percentages describe the distribution of dataset records, rather than unique peptides or population allele frequencies. **(B)** Pearson correlation between predicted residue-level presentation scores and experimentally annotated ligand density for HLA-DRB1*04:01, HLA-DRB1*07:01 and HLA-DRB1*15:01. For each protein, ligand density was defined as the number of distinct annotated peptide spans covering each residue. Prediction scores were obtained by taking the maximum score among all covering peptide windows of 12–25 amino acids. NetMHCIIpan 4.3 EL ranks were sign-inverted so that higher scores consistently indicate stronger predicted presentation. Pearson correlations were calculated separately for each protein and averaged within each held-out test fold. Performance of the MESSI predicted immune risk and NetMHCIIpan 4.3 was evaluated by 5-fold cross-validation for each HLA-DRB1 allele. Both methods were evaluated on identical held-out test proteins in each fold. Metrics were calculated separately for each protein and averaged within each test fold; mean and standard deviations across 5 test folds are plotted. **(C)** Residue-level discrimination of IEDB peptide annotations, measured by the area under the receiver operating characteristic curve (ROC-AUC). Residues covered by at least one annotated ligand were labeled positive, and uncovered residues were treated as negative for this evaluation. Both MESSI and NMP used the same residue-score aggregation and held-out test folds as in B. ROC-AUC was calculated separately for each protein and averaged within each test fold. Bars show the mean ± standard deviation of these fold-level averages across five-fold cross-validation. **(D)** Single-protein inference latency as a function of sequence length for the HLA-DRB1*07:01 MESSI immune risk predictor and NetMHCIIpan 4.3. All three series use the same 14 protein sequences, spanning 59–1,500 amino acids, and evaluate all peptide windows of 12–25 amino acids. CPU measurements were obtained on the same Apple M2 Pro host, with one intra-operation CPU thread for MESSI; GPU measurements used one NVIDIA RTX PRO 6000 Blackwell Server Edition. Points show median latency over 30 MESSI CPU calls, 100 MESSI GPU calls or two independent NetMHCIIpan subprocess calls per protein, after excluding warm-up measurements. MESSI timings include the complete prediction call (sequence encoding, window enumeration and scoring, and residue-level aggregation) but exclude checkpoint loading. NetMHCIIpan timings include subprocess startup and model initialization. The vertical dotted line marks the 1,022-residue chunking boundary. Inference time is shown on a logarithmic scale. **(E)** Illustrative whole-protein presentation-risk landscape for P00352, a 501-amino-acid protein, evaluated for HLA-DRB1*07:01 using the MESSI immune risk predictor and NetMHCIIpan 4.3. The upper track shows IEDB-annotated ligand spans, forming three overlap-connected regions. The middle track shows the continuous residue-level risk field obtained by log-mean-exp aggregation of scores from covering peptide windows, with Gaussian smoothing applied for visualization only (σ = 1.2 residues). The lower track shows NetMHCIIpan 4.3 peptide windows with EL percentile ranks ≤2%; each horizontal segment represents one predicted binding window, and vertical stacking separates overlapping calls rather than encoding prediction strength. All tracks share zero-based residue coordinates.

To benchmark our trained immune risk predictor, we compared it to NetMHCIIpan-4.3 (referred hereafter as NMP), a widely used MHC-II peptide prediction model that has been validated both computationally and experimentally (*33*). Since NMP classification happens on a peptide level, we first tested whether our predictor can accurately determine residue level risk for efficient steering of generative models. Using 5-fold cross validation, we found that MESSI’s immune risk predictor achieved a higher Pearson correlation with ground truth labels than NMP (Fig. 2B), faithfully capturing the residue-level pattern of annotated eluted-ligand coverage (*33*). In the classification task, we found that MESSI’s immune risk estimator achieved a similar area under the receiver operating characteristic curve (ROC-AUC) value compared to NMP across all 3 alleles that we tested (Fig. 2C). Moreover, by training a neural network to directly learn the eluted-ligand data, we found that MESSI’s immune risk predictor was up to 1,500x faster compared to NMP and, unlike NMP, is compatible with GPU for faster inference to better couple with generative models (Fig. 2D). As an example to visualize the prediction by MESSI immune risk prediction head, we zoomed in on human Aldehyde dehydrogenase 1A1, a 501-amino-acid protein, containing a ground truth peptide from IEDB and multiple predicted strong binders by NMP (Fig. 2E). Overall, we see that both NMP and MESSI’s immune risk predictor can accurately predict the ground truth T cell epitopes with minimal false positives (Fig. 2E). This data established the performance of our trained single allele estimators, which we next apply to guide and constrain our diffusion model during sequence generation to avoid T cell epitopes.

With this efficient and accurate immune risk prediction, we set out to test whether it can be used to steer a discrete diffusion model for inverse folding toward lower MHC-II presentation risk. MESSI provides guidance in certain predefined diffusion timesteps: from a selected partial diffusion state, MESSI first performs Lookahead by sampling several complete sequences from the current diffusion state and evaluating their immune risk landscapes (Fig. 1). The endpoint complete sequence with minimal whole-protein-level immune risk and its original partial state jointly inform the next generative state under a Feedback operation: residue choices supported by their contribution to reduced whole-protein-level immune risk in the complete candidate context are carried forward, while positions associated with increased local-residue-level risk are reopened to masked states. These decisions define a feedback-updated partial state at an earlier point in the denoising schedule. Denoising resumes from this state and advances to the next predefined timestep, which allows reopened positions to be resolved in several generation steps under retained sequence information and an evolving context. The resulting partial state on this predefined timestep becomes the source for the next lookahead round. Repeating this lookahead–feedback cycle preserves selected sequence information as an explicit generative memory while allowing earlier choices to be revised as their contexts change. MESSI thus turns complete-sequence immune evaluation into cumulative generative search for structure-compatible sequences with reduced MHC-II presentation risk (see Supplementary Text for model and algorithm details).

To examine how feedback reshaped generation, we constructed a set of proteins with high predicted immune risk, spanning diverse sequences and structural topologies to test the performance of MESSI (fig. S1). We first compared unguided diffusion, post-hoc selection and MESSI with increasing feedback depth on deimmunizing 3 different HLA alleles (Fig. 3A, fig. S2A). The redesigned distributions shifted toward lower immune risk burden validated both by NMP and MESSI risk predictor, with later rounds achieving further reductions across 3 alleles tested (Fig. 3A, fig. S2B). This cumulative shift, observed across the entire set of tested proteins and within individual proteins, highlights that our feedback algorithm progressively and efficiently directs the generation of low-risk sequences, reaching lower levels of immune risk than what can be achieved by post-hoc selection of minimal-risk variants from unconditioned sequence generation (Fig. 3B-C).

**Figure 3.**
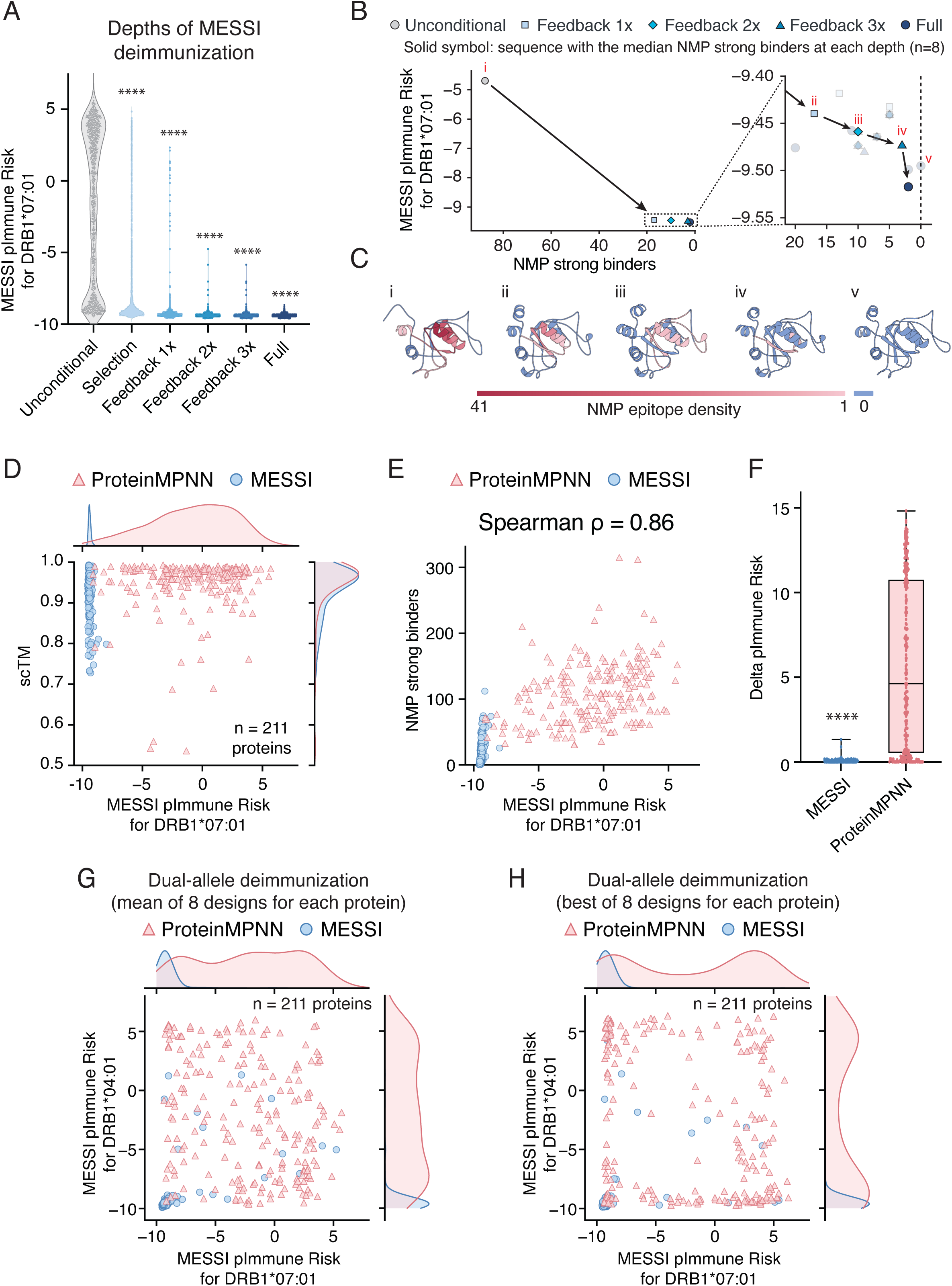
MESSI reduces predicted immune risk while preserving protein structure across HLA alleles. **(A)** A violin plot showing that increasing search intensity shifts the candidate distribution toward lower MESSI pImmune risk (predicted immune risk) across 196 proteins for HLA-DRB1*07:01. Unconditional generation is compared with selection without recursive feedback (D0), cumulative search through depths 1–3, and the complete algorithm (Full). The depth-resolved comparison captures improvement across the candidate population, rather than only in the best sequence recovered for each protein: recursive feedback expands the search pool from which MESSI selects its outputs. Eight candidates per protein are shown at each stage. Search candidates are ranked jointly by MESSI pImmune risk and positive-mass density using Pareto selection. White circles indicate means; boxes indicate interquartile ranges, with medians marked. A one-way ANOVA test is ran between all 6 conditions and multiple comparisons are calculated against unconditional generation. ****, p < 0.0001. **(B)** Search progression for 6SZY_B illustrates how this population-level improvement is realized within an individual protein. Successive feedback stages recover candidates with lower MESSI pImmune risk and fewer NetMHCIIpan4.3-predicted strong-binding windows, culminating in a final algorithm output with no predicted strong binders. **(C)** The five structural examples i-V show an unconditional candidate, immune-ranked representatives from depths 1–3, and the lowest-MESSI-pImmune-risk sequence among the final outputs with zero strong-binding windows. This example connects optimization of MESSI’s learned immune objective to reduced peptide-level burden under an external predictor. Structures were predicted using ESMFold2; strong binders are overlapping 12–25-residue peptide windows with strong binders defined by top 2% as the EL percentile-rank cutoff. **(D)** Predicted immune risk and structural self-consistency of MESSI and ProteinMPNN designs on the HLA-DRB1*07:01 high-risk test set. Both methods were evaluated using 8 final MESSI output sequences and eight ProteinMPNN sequences per target. Each point represents the within-protein mean pImmune risk and mean structural self-consistency across these eight sequences. Immune risk was evaluated using the frozen HLA-DRB1*07:01 pImmune risk from MESSI, with lower scores indicating lower predicted presentation risk. Structural self-consistency was measured by scTM, the TM-score between the ESMFold2-predicted structure of a designed sequence and its target backbone; higher values indicate closer structural agreement. Blue circles and pink triangles denote MESSI and ProteinMPNN, respectively. Marginal curves show the distributions of protein-level mean risk and mean scTM, with each density curve independently normalized to unit peak. **(E)** Association between pImmune risk and NetMHCIIpan-predicted strong-binder counts for the same proteins and output sequences as in D. Each point represents the within-protein mean pImmune risk and mean strong-binder count across eight sequences. Strong binders were defined as overlapping peptide windows of 12–25 amino acids with a NetMHCIIpan-4.3 percentile rank below 2 %. Counts therefore represent predicted strong-binding windows rather than distinct epitopes. Spearman’s correlation was calculated across the pooled protein-level observations from both methods (ρ = 0.86; 422 observations representing 211 targets per method). **(F)** Predicted immune-risk difference between structure-based and immune-based sequence selection within each 8-sequence output panel. For each target protein and method, Δ immune risk was calculated as the pImmune risk of the highest-scTM sequence minus the minimum pImmune risk among the eight sequences. Values near zero indicate that the most structurally self-consistent sequence also has near-minimal immune risk within the panel; larger positive values indicate a greater immune-risk cost of selecting solely for structural self-consistency. Boxplots summarize the 211 protein-level differences per method. Paired t-test (two-sided) is used between the MESSI and proteinMPNN. ****, p < 0.0001. **(G)** Dual-allele immune-aware design on a test set assembled from the HLA-DRB1*07:01 and HLA-DRB1*04:01 high-risk test sets. MESSI jointly targeted the 2 allele-specific immune objectives, and both methods were evaluated on the same 211 target proteins. Each point represents the within-protein mean pImmune risk across the method’s final output sequences, with HLA-DRB1*07:01 risk on the horizontal axis and HLA-DRB1*04:01 risk on the vertical axis. Both allele-specific means were calculated from the same set of sequences. Lower-left positions indicate lower predicted risk for both alleles. Marginal density curves are independently normalized to unit peak. **(H)** Best-immune sequence outcomes for the same dual-allele comparison as in G. For each target protein and method, one sequence was selected by minimizing the joint immune objective over the final output panel. The same joint criterion was applied to MESSI and ProteinMPNN, with ties resolved deterministically by sequence identifier and design index. Each point shows the 2 allele-specific pImmune risk scores of that single selected sequence, rather than independently selected per-allele minima. Both methods are shown for the same 211 target proteins. Marginal density curves are independently normalized to unit peak.

Next, to benchmark MESSI against a widely used and extensively experimentally validated inverse folding model (ProteinMPNN) with post-hoc filtering approaches (*50*), we used both MESSI and ProteinMPNN to design 8 sequences per starting structure, for a total of 211 proteins, and evaluated the Pareto frontier of immune risk with structural recovery for each of the 3 HLA-DR alleles. We found that MESSI is able to significantly deimmunize the set of protein structures while retaining a good structural recovery rate with scTM>0.7 for all test proteins across all 3 tested alleles (Fig. 3D, fig. S3A-B). In particular, we found significantly reduced immune risk distribution for all 3 alleles without sacrificing structural metrics compared to unconstrained ProteinMPNN, which only optimizes for structural recovery, leaving large variance in predicted immune risk both in terms of mean and the best of the 8 designed sequences (Fig. 3D, fig. S3C-D). To make sure that predicted immune risk by MESSI accurately reflects the T cell epitopes burden for diverse proteins, we again evaluated the predicted immune risks against an orthogonal NMP prediction. We found highly correlated rankings with a Spearman correlation of 0.86 between MESSI’s predicted immune risks with strong binders predicted by NMP (Fig. 3E), which reinforces the accuracy of MESSI’s immune risk estimation. Next, we evaluated the tradeoff between structural recovery and reduced immune risk by measuring the difference between the best structural recovery sequence (defined by highest scTM) and the lowest immune risk sequence. As expected, we found MESSI only generates sequences with low immune risk since the diffusion process is constrained by the immune risk module whereas regular inverse folding models like ProteinMPNN have large variance between different proteins with significant tradeoff (Fig. 3F, fig. S4).

Lastly, we tested whether MESSI can be used to deimmunize risks for multiple alleles of interest since for any given protein therapeutics to be immunogenicity free in the clinic, it has to satisfy a minimum of 3 MHC class II molecules that are present in any human. Here as a proof of concept, we constrained MESSI to generate sequences by avoiding immune risks against both DRB1*04:01 and DRB1*07:01 molecules. Similar to single allele design, we found that MESSI is efficient in generating sequences that avoid epitopes for both alleles of interest, whereas unconditional ProteinMPNN had large variance in immune risks across both alleles (Fig. 3G-H). Overall these data support MESSI as an inverse folding model that can deimmunize a protein of interest while efficiently exploring the Pareto frontier of immune risk with structural recovery.

### Deimmunization of luciferase

Self-consistency between the predicted structures and the original input is not a guarantee of protein function. In fact, inverse folding models in general can produce many sequences that are not functional due to various issues, including inaccuracy from structural prediction models and neglect of molecular dynamics. Thus, it is critical to validate that MESSI can indeed design sequences that recapitulate the desired protein function rather than just the predicted structure. On this front, we first applied MESSI to deimmunize LuxSit-i, a small (13.8-kDa) luciferase that was previously designed *de novo* using deep learning-based hallucination (*76*). To maximize the probability of preserving enzyme activity in addition to structure, we restricted the inverse folding process by fixing amino acid residues that are known to be involved in catalysis. Using AlphaFold3-predicted structure of LuxSit-i (*44*), we identified 22 sites with functions ranging from previously identified catalytic active site to residues that line the substrate pocket and involved in packing (Fig. 4A). We then implemented MESSI by designating these 22 residues to be fixed during diffusion and allowed MESSI to generate the rest of the sequences. At the end of generation, we selected 5 sequences based on their predicted structural quality and sequence diversity (Fig. 4B, fig. S5A). These LuxSit-i variants had 53-57% sequence identity relative to the parental LuxSit-i (Fig 4B), but folded into structures nearly identical to the parental protein (Fig. 4C) while having 0 predicted epitopes to the target HLA-DR*15:01 allele (Fig. 4D, fig. S5B). However, it is known that predicted static structure does not guarantee protein function (*35*, *77*–*79*). Thus, to experimentally validate their functions, we synthesized these LuxSit-i variants *in vitro* using a PURExpress cell-free translation kit and measured their luminescence in the presence of diphenylterazine (DTZ) (*76*), a synthetic luciferin previously used for characterizing LuxSit-i (Fig. 4E). We observed that all 5 variants generated by MESSI had detectable luciferase activity, with 1 variant (MESSI-D2) achieving a luminescence output similar to that of parental LuxSit-i (Fig. 4F, fig. S5C). Overall, these results suggest that MESSI can redesign an enzyme to minimize its predicted immunogenicity against a specified human MHC II molecule, while retaining nearly identical structure and function.

**Figure 4.**
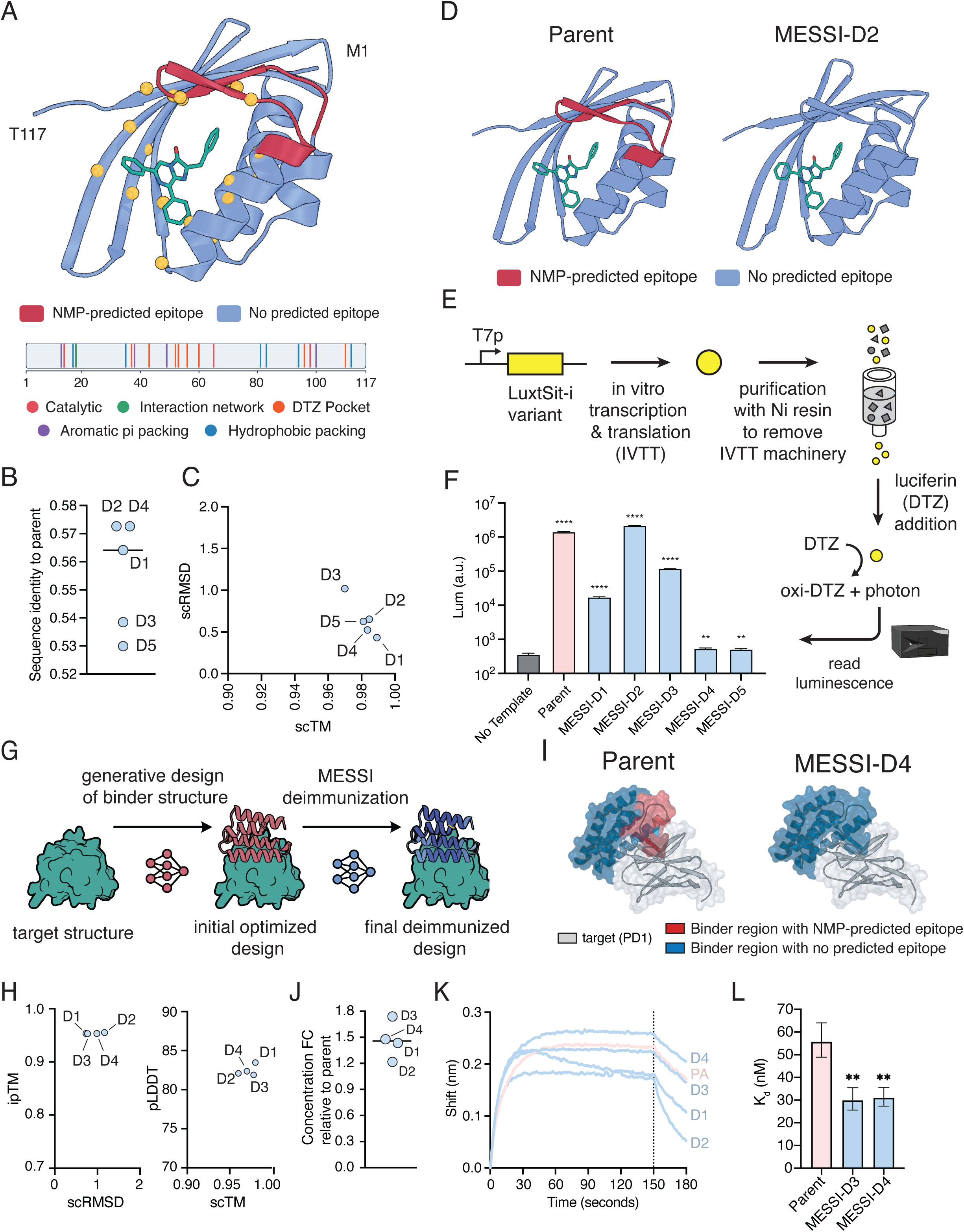
MESSI generates functional, epitope-free variants of a luciferase and an anti-PD-1 binder. **(A)** Fixed amino acid residues for MESSI-based deimmunization of LuxSit-i. The structure of LuxSit-i (shown in blue and red) with its luciferin substrate (shown in green) is generated by Alphafold-3. The color on LuxSit-i denotes the number of overlapping NMP-predicted HLA-DRB1*15:01 epitopes at each amino acid position. Fixed amino acid residues are depicted as yellow-colored spheres. **(B)** Sequence identity of MESSI-generated LuxSit-i variants relative to the parental LuxSit-i. **(C)** Structural metrics of MESSI-generated LuxSit-i variants. **(D)** Comparison of epitope density between parental LuxSit-i and a MESSI-generated variant, MESSI-D2. All structures were generated by AlphaFold-3. Color denotes the number of overlapping NMP-predicted HLA-DRB1*15:01 epitopes at each amino acid position. **(E)** Experimental validation of luminescence from MESSI-generated LuxSit-i variants. Gene fragments encoding the parental and MESSI-designed proteins were synthesized and used as linear DNA template for in vitro RNA transcription and protein translation (IVTT). Afterwards, the reaction product was purified with a nickel spin column to remove the His-tagged IVTT machinery. A final concentration of 25 µM of diphenylterazine (DTZ) was added to the purified proteins. Luminescence was recorded on a plate reader every minute over 2 hr. **(F)** Peak luminescence recorded within 2 hours after DTZ addition. “No template” denotes the negative control condition where no DNA template was added to the IVTT reaction. Error bars, standard deviation. n = 3 replicates. A two sided unpaired t-test with welch’s correction is calculated between no-target condition and each of the MESSI designs and parent. **, p < 0.01; ****, p < 0.0001. **(G)** Computational workflow for designing a deimmunized binder against PD-1 using MESSI coupled with BindCraft. In particular, MESSI is inserted at the end of the original BindCraft workflow to replace the soluble MPNN inverse folding model to design sequences against the optimized structures. **(H)** Structural metrics of MESSI-generated variants of anti-PD-1 binder. **(I)** Comparison of epitope density between the parental anti-PD-1 binder and a MESSI-generated variant, MESSI-D4. All structures were generated by AlphaFold-3. Blue colors denote a binder region that does not contain NMP predicted epitopes. Red color denotes binder regions with NMP-predicted HLA-DRB1*15:01 epitopes greater than 0. White color denotes the PD-1, the target protein for the designed binder. **(J)** Strep-tagged minibinders are produced with Genscript IVTT kit. After the production, the binders are purified with streptavidin beads and eluted in 1 mL of PBS. The dot plots show the concentration (expression) fold-change (FC) of the MESSI-generated binders relative to the parental binder designed by BindCraft. **(K)** Bio-Layer Interferometry (BLI) binding curves for measuring the binding parameters of MESSI-designed anti-PD-1 binders against the target PD-1 protein. The assay was performed at 30 °C and at 1,000 rpm. His-tagged PD-1 protein was firstly immobilized onto HIS1K biosensor on Octet RED384. Binders was applied at a concentration of 1 µM as an analyte for association and dissociation steps for 150 seconds and 30 seconds respectively. PA denotes the parent binder. **(L)** Equilibrium dissociation constant (K_d_) of anti-PD-1 binder variants. K_d_ was derived from the association (K_a_) and dissociation (K_dis_) rate constants measured from the five concentration scan BLI curves. Error bars represent the 95% confidence interval of the fitted K_d_. Differences in K_d_ were assessed by a two-sided z-test using moving-block bootstrap standard errors of the global fit (85 seconds blocks, 1,000 resamples; Holm-corrected for comparisons). **, p<0.01

### Binder design with immunogenicity awareness

Recently, a wave of generative models has been developed to design protein binders against a protein of interest (*24*, *26*, *27*, *29*, *80*). This is a therapeutically important goal, as the ability to design proteins to bind to a specified epitope without extensive experimental screening would revolutionize our ability to design macromolecule medicines for treating various diseases. However, this class of models in general suffers from immunogenicity liabilities of the sequences they generate. These sequences generated *in vitro* would not simulate or go through the thymic selection to be tolerized in humans and the training input for these models in general contain mostly non-human sequences (*41*, *81*). Thus, *de novo* designed minibinders in general would be expected to have strong immunogenicity in humans. Even in cases of scFv and antibody design, where only the complementarity-determining region is opened up for design and human framework region is preserved to maximize “humanness score”, there is still a possibility of immunogenic reaction due to the complexity of MHC class II processing and presentation. In fact, even for antibody sequences generated from humanized mice models or other animal immunization campaigns, there have been reports of clinical immunogenicity, highlighting the importance of deimmunization for antibody based protein therapeutics in general (*23*, *82*, *83*).

For proof of concept, we decided to couple MESSI with BindCraft (*27*), a well validated binder generation pipeline to determine whether we could successfully deimmunize *de novo* designed minibinders. We chose to focus on a validated binder against Programmed cell death protein 1 (PD-1), a core protein in the T-cell immune checkpoint pathway (*84*). Blockade of the PD-1/PD-L1 pathway with an inhibitory binder is the foundation of modern cancer immunotherapy (*1*). BindCraft works through Alphafold 2 based hallucination to arrive at an optimized structure against the target protein and then uses soluble ProteinMPNN to perform inverse folding to design sequences for the desired binder structures (*85*). Here, we inserted MESSI to replace the ProteinMPNN inverse folding model used in this pipeline to see if we can steer the inverse folding sequence generation part to be immunogenicity aware (Fig. 4G). We again fixed 57 contact hotspot residues similar to the BindCraft protocol and allowed MESSI to only design the remaining positions in the binder backbone. We selected 4 generated sequences all with 0 predicted epitopes for human HLA-DRB1*15:01 allele with a median sequence identity of 81% to the parent sequence generated by BindCraft (Fig. 4I, fig. S5D). Although computational metrics are known to be not reflective of binding strength (dissociation constant K_d_), they still provide a computational filter for judging whether these binders will likely bind in the predicted orientation and epitope (*44*). We found a high structural confidence recovery rate for the MESSI designs (Fig. 4H) with iPTM scores above 0.9 for all 4 binders, each of which was well above the typically used cutoff for a positive binder (*27*). We proceeded to purify these 4 designed binders using a C-terminal strep tag for purification, and found that all 4 designed binders expressed at higher concentrations than the parent sequence in the cell free E.coli expression system (Fig. 4J). Interestingly, the model without explicitly training for stability or solubility can generate designs with higher expressability than their parent in this small PD-1 binder campaign, likely hinting the implicit prior learned through the diffusion language mode backbone (*48*). Lastly, we performed Bio-Layer Interferometry measurement to characterize the binding affinity of these binders to the target PD-1 protein first at a fixed concentration of 1 µM and found that all 4 binders had detectable binding to the target protein (Fig. 4K). Since a single concentration BLI measurement can not accurately determine K_d_, we measured the binding kinetics of the MESSI-D3, MESSI-D4, and the parent binder using 5 concentrations from 12.5 nM to 200 nM with 300 seconds of dissociation time (fig. S5E). With this data, we estimated K_d_ and found that MESSI-D3 and MESSI-D4 had significantly better binding affinity than the original parent sequence with MESSI-D3 reaching a K_d_ of 30 nM (Fig. 4L). This data along with the luciferase data support MESSI as a generalizable inverse folding model that can be applied to various proteins to minimize their predicted MHC-II epitopes, while retaining structure and function.

### Deimmunization of Uricase

Lastly, we applied MESSI to deimmunize a clinically important protein drug urate oxidase (uricase) (*86*, *87*). During evolution, humans have lost the functional uricase gene, which in other species encodes an enzyme that breaks down insoluble uric acid into a soluble compound called allantoin that can be easily excreted by the body. Consequently, when uric acid levels are elevated in our system (hyperuricemia), it has severe consequences including sharp crystals in joints, as seen in chronic gout patients, as well as kidney stones and potential kidney failure (*88*). Thus, it would be ideal if we could design therapies that can durably lower uric acid with minimal side effects. In light of the lost functional uricase gene in our genome, functional uricases from other species are of significant interest because evolution has already provided templates for this specific function. Indeed, there are two FDA approved uricase products in the clinic. *Aspergillus flavus* derived uricase rasburicase is used to control hyperuricemia associated with tumor lysis (*86*), and a PEGylated porcine–baboon chimeric uricase is used in the treatment of refractory gout (*87*). However, immunogenicity remains a limitation even for these approved uricase therapies: high-titer antibodies have been detected against uricase and they are associated with accelerated drug clearance, loss of sustained urate lowering, and increased risks of infusion reactions (*12*, *13*). These responses include antibodies against PEG as well as the uricase component, and due to these constraints previous efforts have also attempted deimmunization with simpler alignment methods (*89*). Here, we leverage MESSI to perform deimmunization of this class of enzymes for safer and more effective use in humans.

We first performed a comprehensive search for uricases across the tree of life, uncovering 6,801 unique sequences. We performed clustering to remove sequence redundancies and then constructed a phylogenetic tree, yielding 3,034 sequences spanning more than 20 phyla (Fig. 5A). Although not every uricase has been validated to be functional, this tree highlights the evolutionary diversity of uricase and represents a large pool of starting points for deimmunization since they yield different levels of potential epitopes (fig. S6A). Moreover, the epitopes fall into various locations throughout the protein, including positions at or nearby catalytic active sites; these epitopes are inherently harder to remove because mutating these positions may compromise enzyme function. We sampled 11 uricase orthologs from this phylogenetic tree to evaluate redesign options across divergent sequence backgrounds, including bacterial, fungal, plant, and animal enzymes (Fig. 5A). The panel included *A. flavus*, *Sus scrofa*, and *Papio hamadryas*, representing source species relevant to clinically used uricases, together with more distantly related orthologs. This sampling allowed us to examine whether the relationship between predicted epitope reduction and functional retention depended on the starting sequence. For each starting sequence, we first fixed 9 previously labelled functionally important sites including the conserved KTH catalytic triad and 6 binding sites for urate and 5-hydroxyisourate (*90*). These 9 sites are also conserved across most of the phylogenetic tree (fig. S6B). Uricase functions as a tetramer whose substrate-binding pockets are formed jointly by neighboring subunits, making intersubunit assembly integral to its catalytic architecture (*90*, *91*). To preserve this organization, we fixed residues forming consistent contacts at the catalytic and tetramer-assembly interfaces across five Protenix-v2 models per parent. In addition to fixing these sites, previous attempts using inverse folding models have revealed the importance of enzyme residues that have high covariance signal during evolution (*47*, *79*). Thus, we deployed a more generalizable conservation and covariance analysis using EVcouplings to scan these sequences and fixed positions with ≥80% conservation or robust covariance scores above the 80th percentile (*92*). With these constraints in mind, we then opened up the rest of the predicted high immune risk regions for re-design by MESSI and sampled up to 3 designs per starting uricase by ranking their monomer structural confidence score, tetramer interface score and urate docking score by refolding the designed sequences using ESMFold2 and protenix-v2 (*25*, *30*).

**Figure 5.**
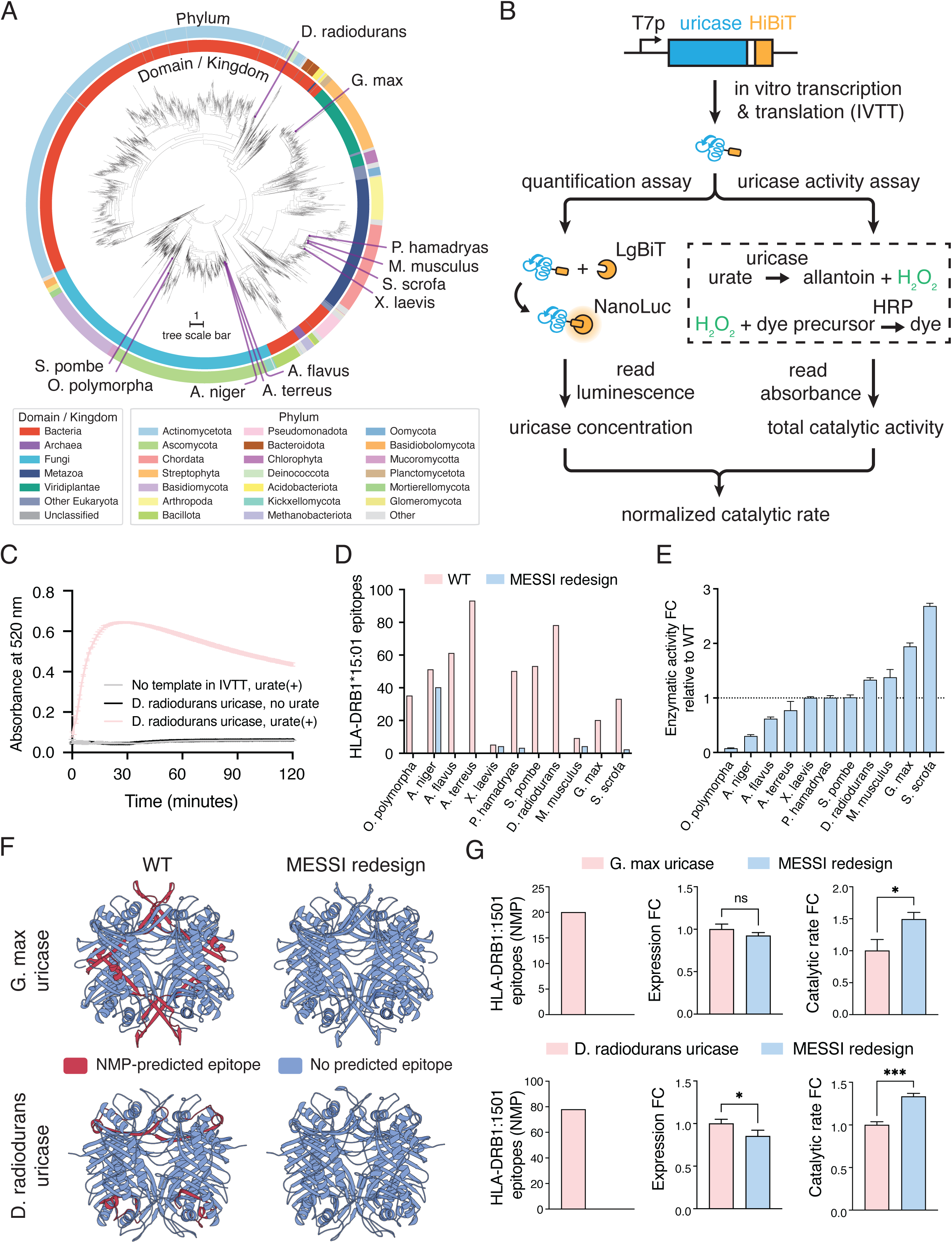
MESSI generates functional, epitope-free variants of tetrameric uricases. **(A)** Phylogenetic tree of uricases. The inner and outer rings show the domain or kingdom, and the phylum, of the organism from which the uricase was identified. Uricases that are deimmunized by MESSI in this work are marked around the tree. **(B)** Experimental assays to quantify enzyme-concentration-normalized catalytic rate from uricase variants. Linear DNA templates encoding wild-type and MESSI-designed uricases were appended with a HiBiT-tag at the C-terminus, and ordered from a commercial DNA synthesis company. Uricases were synthesized with a cell-free in vitro transcription and translation (IVTT) kit. Synthesized uricases underwent a quantification assay, where addition of the HiBiT-complementary LgBiT protein leads to reconstitution of the NanoLuc luciferase, allowing uricase concentration to be measured based on luminescence reading. In parallel, synthesized uricases underwent a uricase activity assay, where the conversion of urate by uricase is coupled to the generation of a dye, allowing measurement of total uricase activity based on the initial slope of absorbance reading at 520 nm. The total uricase activity is normalized by the enzyme concentration obtained from the quantification assay, resulting in the intrinsic catalytic rate of the uricase. **(C)** Representative absorbance curves from the uricase activity assay. Gray and black curves represent two negative control conditions, either with no DNA template at the IVTT step, or with no urate in the dye-forming reaction, respectively. Error bars, standard deviation. n = 3 replicates. **(D)** Number of NMP-predicted HLA-DRB1*15:01 epitopes in WT uricases (pink) versus their MESSI-redesigned variants (blue). For each WT uricase, the variant with the minimal number of epitopes was shown. X-axis, the organism where the wild-type uricase comes from. **(E)** Catalytic rate of the MESSI-redesigned uricases, expressed as FC compared to their wild-type parental proteins. For each wild-type uricase, up to 3 MESSI-redesigned variants were synthesized and tested; the variant with the highest catalytic rate was shown. X-axis, organism where the wild-type uricase comes from. Error bars, standard deviation. n = 3 replicates. **(F)** Comparison of epitope density between the WT uricase and their MESSI-redesigned variant, for two representative uricases. Uricases shown are in their tetrameric form. All structures were generated by Protenix-v2. Color denotes the number of overlapping NMP-predicted HLA-DRB1*15:01 epitopes at each amino acid position. **(G)** Epitope count, expression and catalytic activity of the WT uricases (denoted in pink) and their MESSI-redesigned variants (denoted in blue) shown in Panel F. Expression was measured with the quantification assay shown in Panel B. Catalytic rate was measured by normalizing the total uricase activity by the enzyme expression, using the workflow depicted in Panel B. All FCs were relative to the parental wild-type uricase. Error bars, standard deviation. n = 3 replicates. Two sided unpaired t-test with Welch’s correction is used. ns, p > 0.05; *, p < 0.05; ***, p < 0.001.

To experimentally test the re-designed uricases, we applied an *in vitro* transcription and translation (IVTT) workflow similar to what we used for the validation of LuxSit-i variants. To separate catalytic activity from expression, we appended a HiBit tag onto the C-terminal of each designed and wild-type (WT) uricase sequence (Fig. 5B). Using this approach, the concentration of the expressed uricase can be quantified based on luminescence following complementation with LgBiT. In parallel, total uricase activity was measured using a Trinder assay, in which urate-dependent hydrogen peroxide production was coupled to horseradish peroxidase–mediated dye formation and monitored by absorbance at 520 nm (*93*). To validate this assay, we chose the WT *Deinococcus radiodurans* uricase and its reaction produced a pronounced substrate-dependent signal, whereas the absorbance from reactions lacking either urate or the IVTT DNA template encoding the uricase remained near background (Fig. 5C). Total activity was estimated by taking the slope of the A520 increases in the initial linear region; normalized activity was calculated by dividing the total activity over HiBiT-measured protein concentration. After re-design, we first used NMP to calculate the epitope burden for HLA-DRB1*15:01 (*33*). Overall, the 11 WT uricases differed substantially in their predicted epitope counts, providing starting points with distinct burdens for the same HLA allele. Across the panel, MESSI re-designs reduced these counts, although the magnitude of reduction varied among orthologs (Fig. 5D) likely due to differences in epitope locations and availability of a functional amino acid substitution. Encouragingly, for some uricases including *G. max* and *D. radiodurans*, we were able to identify designs that have 0 predicted epitopes. Next we tested whether the re-designed variants and their parents have detectable uricase activity. To high throughput test these variants, we assayed the enzyme activity by inputting the same amount of IVTT reaction into our Trinder assay and used the Hibit signal to normalize for expression differences. We found that all 11 WT uricases were active and for 7 of the 11 starting points, we found MESSI redesigns that exhibited enzymatic activities close to or above their corresponding wild types (Fig. 5E). For these variants, 4/11 had higher expression than their WT counterparts, and all re-designs had detectable expression within 50% of the WT levels (fig. S6C). In particular, the best *S. scrofa* redesign was around 2.7-fold more active than the original parent, showing that the diffusion language model based inverse folding model can learn substitutions that improve activity similar to the use of the ESM-IF to improve antibodies and other proteins (*34*, *45*, *47*).

Lastly, we zoomed into two re-designed sequences with 0 predicted epitopes (*G. max* and *D. radiodurans* designs) from highly active WT uricases, which illustrate the simultaneous reduction of predicted epitope burden and retention of biochemical function (Fig. 5F-G). We observed that MESSI can re-design enzymes with similar expression level and biochemical activity while avoiding entirely predicted epitopes against the target HLA-DRB1*15:01 allele. Overall, the 3 examples shown here establishes MESSI as a generalizable platform to simultaneously achieve deimmunization and maintain or improve function for various proteins of interest.

## Discussion

Here, we presented a novel generative model to perform protein deimmunization that is more efficient than the currently available tools in terms of both the levels of deimmunization and retention of structure and function. We systematically validated MESSI across 3 different proteins including luciferase, minibinder and uricase, which span diverse protein topologies, multimeric states, and catalytic functions. Encouragingly, we found that MESSI is efficient in finding the optimal Pareto frontier between T cell epitopes and protein function. For both single allele and multi-alleles (2 alleles shown here), MESSI can remove all predicted epitopes without sacrificing, and sometimes even improving, the protein’s function. The ability to optimize sequence against multiple HLA alleles computationally could contribute to off-the-shelf deimmunized protein therapeutics that can be applied over a genetically diverse patient population. In this work, we exclusively focused on HLA-DRB1 molecules due to the abundance of eluted ligand dataset for this particular subset. However, to truly deimmunize a protein for any given person, the model has to delete all epitopes for a minimum of 3 different molecules by considering HLA-DR, DP and DQ proteins. From a population perspective, the numbers quickly scale up to more than 20 alleles of interest in order for any protein drug to remain immune risk free for a genetically diverse patient group. Given the capabilities of the current model, simultaneously satisfying more than 20 alleles might still be challenging for all proteins of interest. With that in mind, models like MESSI could allow customized protein drugs to be generated for a particular patient by specifically designing against the patient’s HLA-allele, which can be easily verified by genome sequencing. More importantly, the probabilistic nature of inverse folding models enables the design of diverse sequences that fold into the same structure and perform similar functions, which allows off-the-shelf reservoirs of multiple drug sequences to be administered in series if they do not share the same epitope to prolong the duration of this particular drug candidate. This is especially attractive for drugs like uricase with thousands of starting structures in evolution to design against.

There are also significant opportunities to improve the current generation of the model by looking at this multi-objective optimization problem from several angles. First, MESSI needs an accurate predictor of the T cell epitope risk. Currently, this relies on training a neural network using the publicly available immunopeptide dataset which is biased against certain alleles. Scaling of this database to cover more alleles, and use of multiple alleles to train the risk predictor could significantly improve the prediction of the epitope risk landscape. In addition, not all presented epitopes will trigger a CD4^+^ T cell reaction, as some presented epitopes might be sufficiently similar to self-peptide and are tolerized in humans. Thus, models to predict T cell receptor vs pMHC interaction and the ability to separate self vs non-self peptide will shrink the number of epitopes to avoid during generation and further increase the likelihood of successful designs. Future works to incorporate other variables that impact clinical immunogenicity including B cell epitopes and protein aggregation propensity into the design stage and testing of these designed sequences in preclinical immunogenicity assays will enable models like MESSI to have a greater impact in the clinic.

## Acknowledgements

We would like to thank the Myhrvold lab for help with cloning, protein expression and instrumentation support in this study; T. Chen, D. Chen, Z. Liu, and J. Wang for computational discussions. We thank H.Y. Chang, C. Tinberg, C. King, R. Potts, Y. Kang, C. Brangwynne, J. Toettcher, and Z. Tang for support and helpful discussions. We thank the Modal academic compute program for supporting this work. We are pleased to acknowledge that the work reported in this paper was performed using the Princeton Research Computing resources at Princeton University. Princeton Research Computing is a consortium of groups including the Princeton Institute for Computational Science and Engineering (PICSciE) and Research Computing at Princeton University. We thank the Princeton Research computing team, Princeton language initiative and Princeton AI for accelerated invention (AI2) initiative for their support.

## Funding

K.J. is supported by Princeton AI Lab Seed Grant, Princeton SEAS Innovation Fund, SEAS seed fund and Omenn-Darling Bioengineering Institute. C.M. is supported by a Princeton AI Lab Seed Grant and by the Princeton Catalysis Initiative. L.N. is supported by the Omenn-Darling Bioengineering Institute Innovators Fellowship and the Charles H. Revson Foundation Senior Fellowship in Biomedical Science.

## Authors contributions

Conceptualization: K.J., Z.C.; Methodology: K.J., Z.C., C.J.Y., C.Z.; Investigation: K.J., Z.C., C.J.Y., L.T.N.; Visualization: X.J., Z.C., K.J., C.J.Y.; Funding acquisition: K.J.; Project administration: K.J.; Supervision: K.J.; Writing: all authors

## Competing interests

Princeton University has filed a US provisional patent on this work (No. 64/154,263) with K.J. and Z.C. as named inventors. K.J. is a co-founder of Foundry BioSciences. All other authors declare no competing interests. No companies are currently involved in commercializing this technology.

## Data and materials availability

Models and codes are available at the following Github repository: https://github.com/JiangLabPrinceton/MESSI

## Figure Captions

**Supplementary Fig 1.**
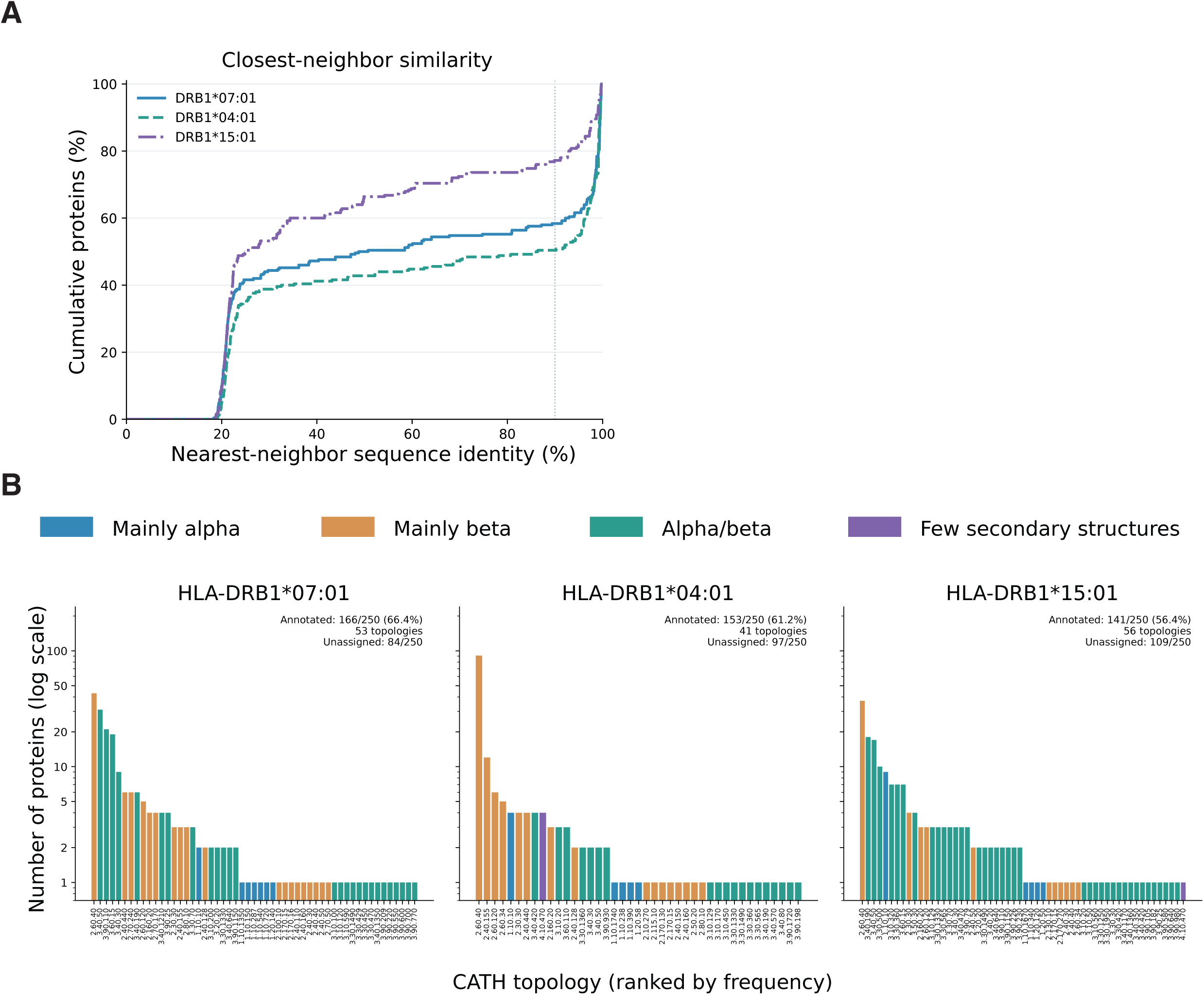
Characterization of the high immune risk protein set. (A) Cumulative distributions of each protein’s highest sequence identity to another protein in the same protein set. Identity was calculated by global BLOSUM62 alignment (gap opening −10, extension −0.5), including gaps in the denominator. Colors and line styles distinguish alleles; the dotted line marks 90% identity. (B) Protein counts per CATH v4.4.0 topology, ranked by frequency within each allele and shown on a logarithmic scale; colors indicate structural class. Exact source-chain assignments required ≥80% domain retention, relative to the larger of the CATH-listed length and source-structure C-alpha count within domain boundaries. Each protein was counted once per topology and could contribute to multiple topologies. Insets report annotation coverage, topology counts, and unassigned proteins; coverage uses all inputs as the denominator.

**Supplementary Fig 2.**
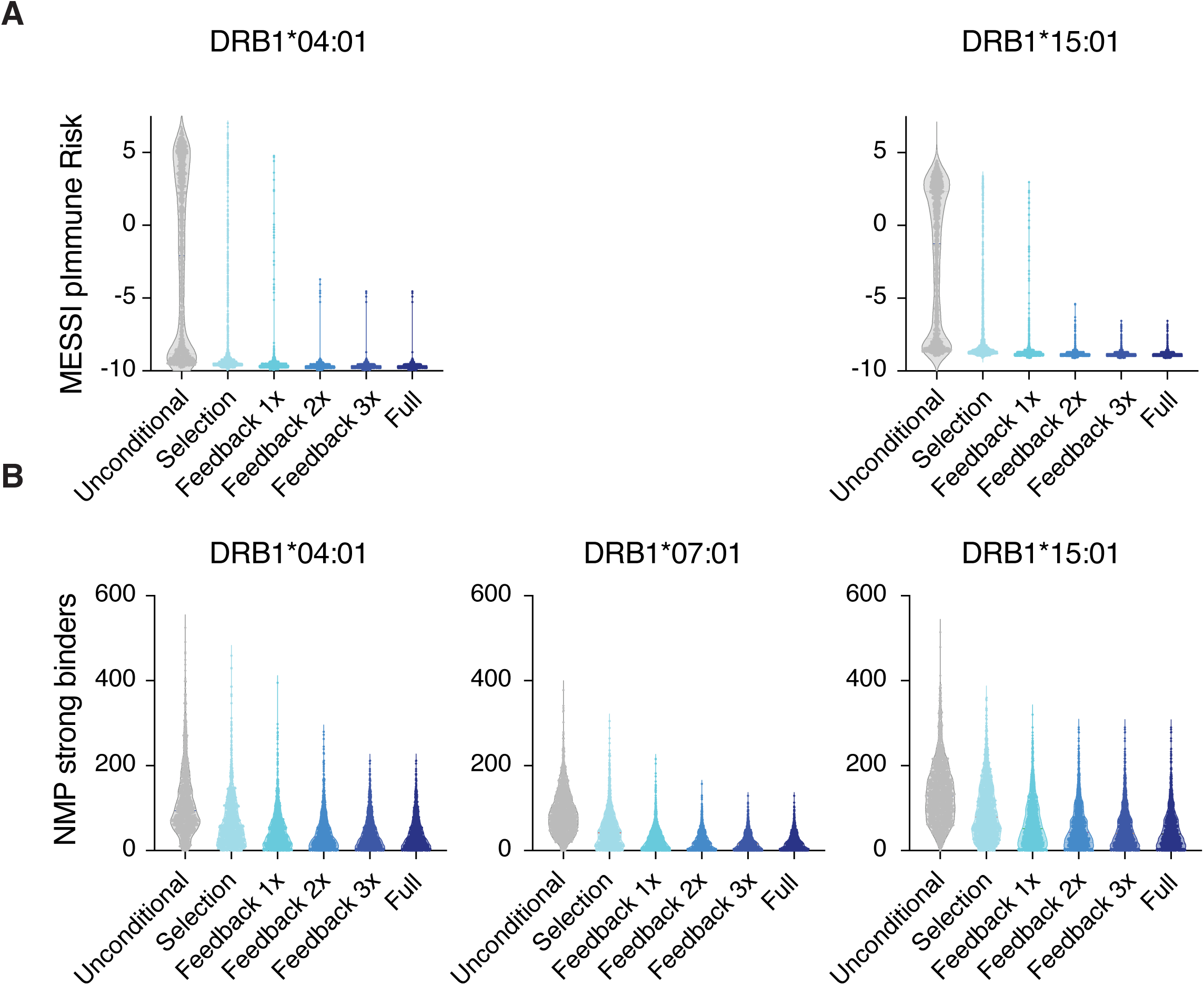
Ablation and mechanistic analysis of MESSI. Violin plots showing that different feedback depths of MESSI ware compared with either unconditional design or post-hoc filtering based on the unconditional design for separate deimmunization campaigns on (A) HLA-DRB1*04:01 and HLA-DRB1*15:01 respectively in terms of the level of deimmunization, calculated by the MESSI immune risk predictor and (B) for HLA-DRB1*04:01, HLA-DRB1*07:01, and HLA-DRB1*15:01 calculated by NMP predicted strong binders.

**Supplementary Fig 3.**
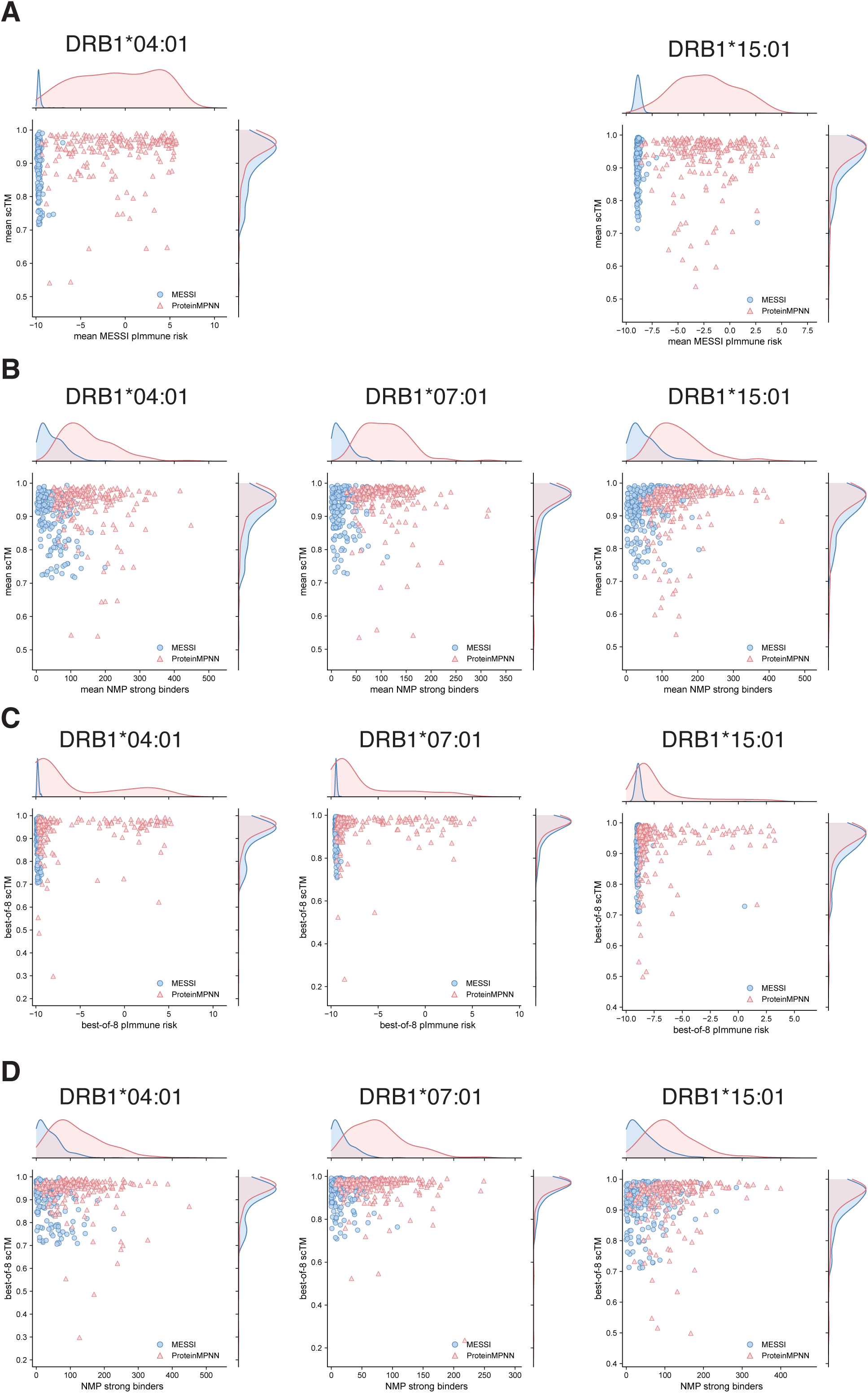
Benchmarking MESSI with ProteinMPNN. MESSI and ProteinMPNN were used to deimmunize the constructed test set by giving a computational budget for 8 sequences per structure. (A) shows the tradeoff between structural recovery (scTM) and predicted immune risk (by MESSI immune risk predictor) for the mean of the 8 designed sequences. (B) shows the tradeoff between structural recovery (scTM) and number of predicted strong binders by NMP for the mean of the 8 designed sequences. (C) shows the tradeoff between structural recovery (scTM) and predicted immune risk (by MESSI immune risk predictor) for the best of the 8 designed sequences in terms of predicted immune risk. (D) shows the tradeoff between structural recovery (scTM) and number of predicted strong binders by NMP for the best of the 8 designed sequences in terms of number of strong binders.

**Supplementary Fig 4.**
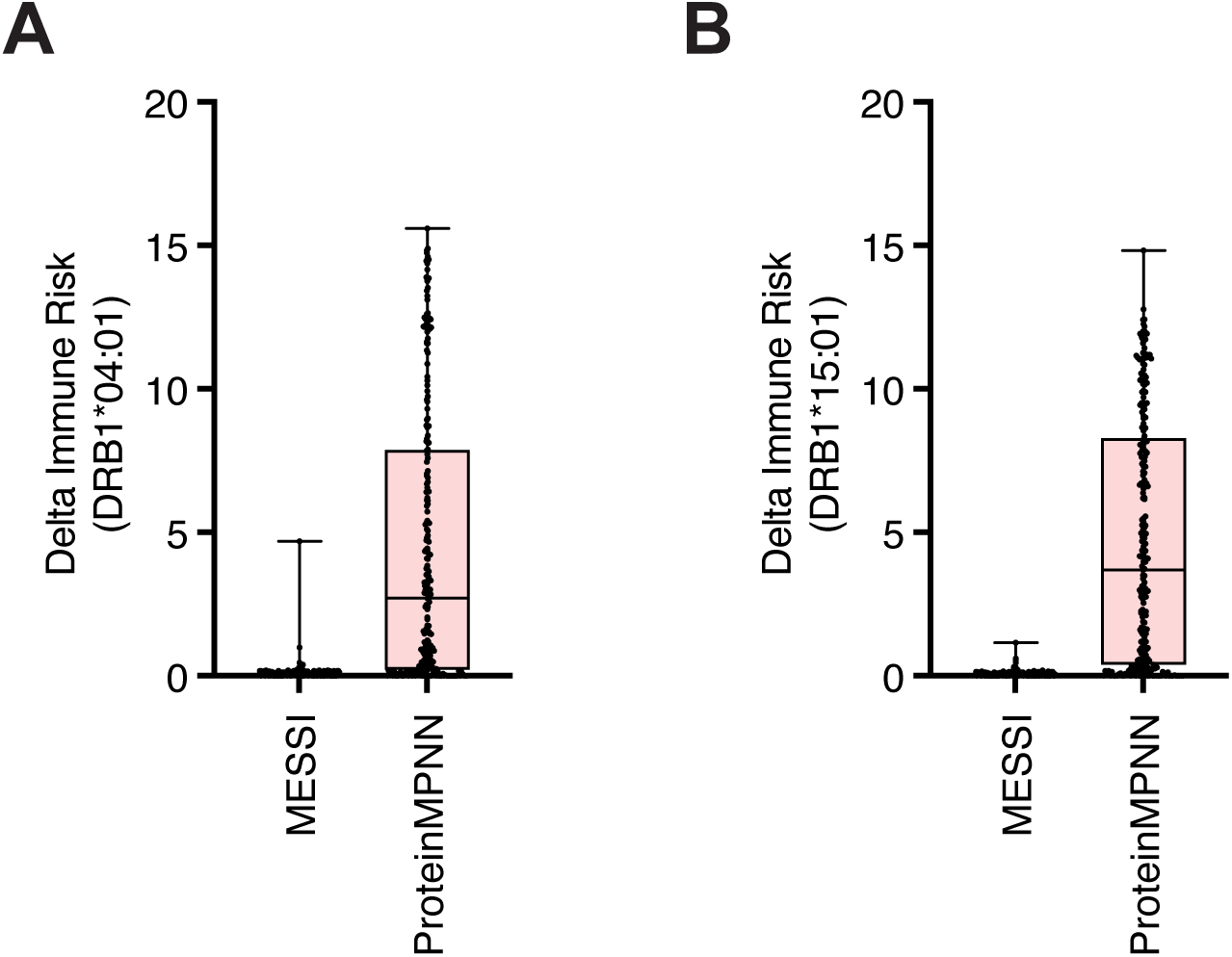
Tradeoff of immune risks with structural recovery. A boxplot with each datapoint showing for each of the tested structures, delta immune risk is calculated by the immune risk of the best structural recovery candidate sequence within the 8 designed sequence minus that of the lowest immune risk candidate within the 8 designed sequences. (A) shows the delta immune risk for MESSI and ProteinMPNN on the 04:01 design campaign and (B) shows that for the 15:01 design campaign.

**Supplementary Fig 5.**
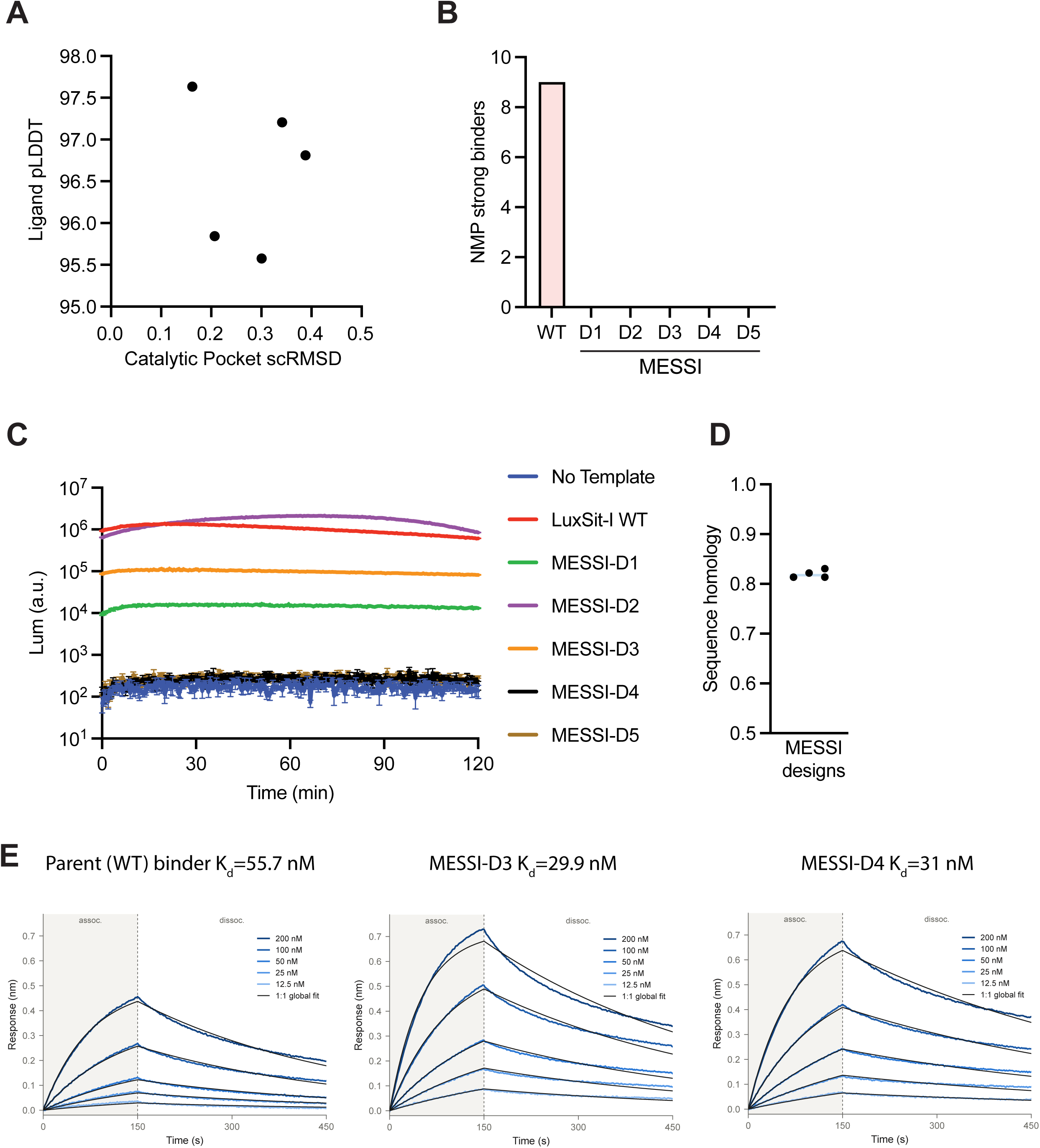
Characterization of the deimmunized luciferase and PD1 minibinder. (A) predicted ligand pLDDT from protenix-v2 predicted structures versus the catalytic pocket (defined as <4 angstroms from the ligand DTZ) self-consistent RMSD for the 5 MESSI redesign variants. (B) Number of NMP predicted strong binders for HLA-DRB1*15:01 allele for the parent luxsit-i luciferase and the 5 MESSI variants. (C) A final concentration of 25 µM of DTZ was added to the purified luxist-i and MESSI designed variants. Luminescence was recorded on a plate reader every minute over 2 hours and the full kinetic traces are shown on a log10 scale. (D) Sequence identity of the 4 MESSI designed PD-1 minibinders relative to the parental sequence from BindCraft. (E) Sensorgrams of five concentrations (12.5-200 nM) of the 2 MESSI designed binders and the parent binder are measured via BLI against immobilized PD-1 antigen with 150 seconds of association time and 300 seconds of dissociation time.

**Supplementary Fig 6.**
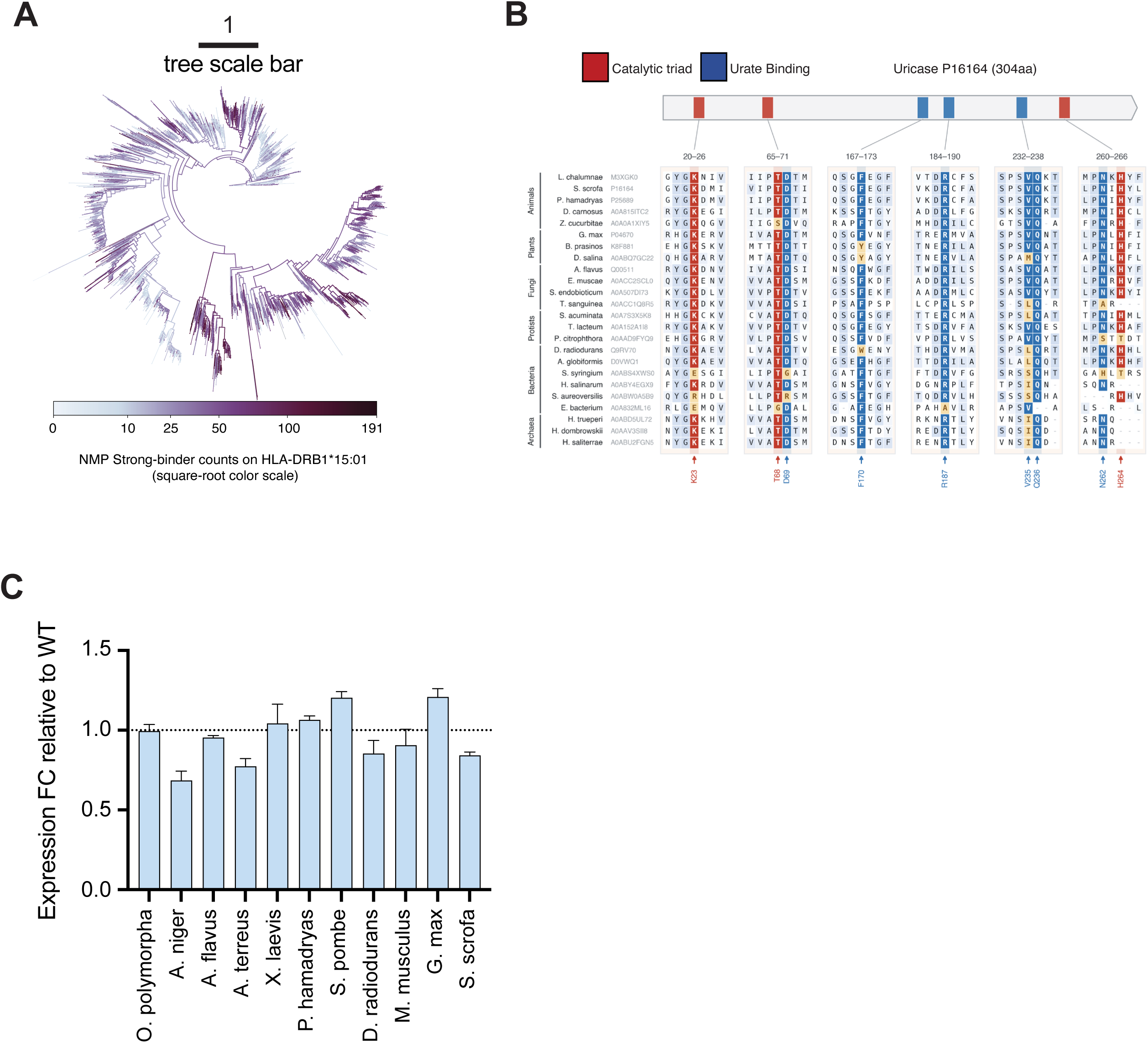
Characterization of the deimmunized uricase. (A) phylogenetic tree of the uricase colored by number of strong binders predicted by NMP on HLA-DRB1*15:01. (B) A representative alignment of the uricase sequences from diverse phylum and kingdom. The red columns highlight the catalytic triad and the blue columns highlight the known urate and reaction intermediate binding sites. (C) Hibit tagged uricase was measured via Promega hibit assay and expression fold change was calculated by dividing the luminescence value of MESSI designed variant over the WT uricase for that design.

## Materials and Methods

### Minibinder synthesis, expression, and purification

Synthetic genes encoding each minibinder were codon-optimized, synthesized, and expressed by GenScript (Piscataway, NJ, USA). All constructs carried a C-terminal Strep-tag II and were produced in a cell-free expression system at a 1 mL reaction scale. Protein was captured in a single step on a Strep-Tactin affinity column and buffer-exchanged into phosphate-buffered saline (PBS), pH 7.2. Purified proteins were stored at -80 °C prior to binding analysis. Recombinant human PD-1 antigen bearing a C-terminal polyhistidine tag was purchased from GenScript (#Z03424), with a reported molecular weight of 32.5 kDa, and supplied at 0.1 mg/mL in PBS. Purity of each preparation was assessed by SDS-PAGE under reducing conditions. Either 2 µg of protein or the maximum volume permitted by the stock concentration was loaded per lane, alongside a molecular weight marker and 2.0 µg bovine serum albumin as a reference standard. Stain-free gels were used and band intensities quantified by densitometry to estimate purity.

### Biolayer interferometry

Binding kinetics were measured by biolayer interferometry (BLI) on an Octet RED384 instrument (Sartorius, Cat. No. FB-20493) using anti-penta-HIS (HIS1K) biosensors (Sartorius, Cat. No. 18-5120) and tilted-bottom 384-well microplates (Sartorius, Cat. No. 18-5166). All steps were performed at 30 °C with orbital shaking at 1,000 rpm, in an assay buffer of PBS, pH 7.2, supplemented with 0.03% (v/v) Tween-20. HIS1K biosensors were pre-hydrated and equilibrated to a stable baseline. 10 μg/mL of his-tagged PD-1 was then immobilized on the biosensor surface. Loaded biosensors were dipped into wells containing Strep-tagged minibinder analyte (1 µM) for a 150-second association step, then returned to the assay buffer for a 30-second dissociation step. Sensorgrams were processed in Octet Analysis Studio v12.2 (Sartorius). Traces were fitted locally to a 1:1 Langmuir binding model to yield association rate constants, dissociation rate constants, and equilibrium dissociation constants. To calculate estimated binding affinity, we took the top 2 binders along with the WT to perform a 5-concentration series scanning. Ligand PD-1 was again immobilized at 10 µg/ml concentration, and then analytes ranging from 12.5-200 nM were applied with 150 seconds of association time and 300 seconds of dissociation time. Sensorgrams were processed in Octet Analysis Studio v12.2 (Sartorius). K_d_ was obtained from global 1:1 fitting of five analyte concentrations (12.5–200 nM), with shared kon and koff. Error bars show the 95% confidence interval of the fit.

### NetMHCIIpan 4.3 prediction of strong binders

For orthogonal validation of immune risk by NMP, we used NetMHCIIpan4.3 by scanning 12- to 25- mer peptides across the entire protein. After scanning using the eluted ligand (EL) mode with the option to predict with context encoding, we selected the top 2% of the EL percentile-rank cutoff as NMP strong binders. This threshold is used throughout the study.

### Constraint generation on uricases

To identify the proper evolutionary and structural constraints for designing uricases, query-centered multiple-sequence alignments were constructed separately for each parental uricase and used to fit PLMC Potts models. Conservation was evaluated after applying sequence-coverage thresholds of 0.6, 0.7 and 0.8. Positions were protected when both gap-excluded conservation and WT residue frequency were ≥0.8 across all three alignments, with a maximum gap fraction of 0.2 (C80). Robust coupling scores were defined as the minimum positional sigma percentile across the top-0.5L, top-L and top-2L coupling sets, where L is sequence length. Positions at or above the 80th percentile were protected only when N_eff/L ≥10 and top-L long-range coupling precision against the five-model tetramer contact consensus was ≥0.60. Five Protenix tetramer models were generated per parent. Chain residues were mapped to the parental sequence. Catalytic interfaces were assigned using bidirectional interchain contacts (<5 Å between heavy atoms) between homolog-mapped Q00511 donor-side residues (Lys11, Thr58 and Asp59) and acceptor-side residues (Phe160, Arg177, Gln229, Asn255 and His257). A unique catalytic chain pairing supported in both symmetry-related interface copies was required in every model. Of the two remaining pairings, the pairing with the larger mean buried surface area was assigned as the assembly interface. Parents with incomplete mappings or unresolved interface assignments were excluded. Residues contacting the opposing chain in both interface copies in all five models were protected. The structural mask was the union of catalytic- and assembly-interface masks; diagonal contacts were excluded. Homolog-mapped functional positions, including the Q00511 Val228-equivalent position, were additionally protected. Interface alanine scanning used a representative tetramer model and the Rosetta ref2015 score function without bound- or unbound-state repacking. ΔΔG_bind was defined as a mutant minus WT binding score in Rosetta energy units (REU). Scores were averaged across the two chain sides within each evaluated interface, and the maximum interface-specific mean was used per position. Positions with ΔΔG_bind ≥1 REU, together with scanned native Ala/Gly/Pro and disulfide-bonded Cys positions, were protected. The union of these masks defined the immutable WT positions in each parent-specific constraint manifest. All protected positions retained their parental amino-acid identities throughout sequence design.

### General procedure for synthesizing enzymes for activity assays

Protein sequences were codon optimized with Twist Codon Optimization for *E. coli* expression. Codon optimized sequences were appended with a constant 5’ sequence containing a T7 promoter and a ribosome binding site, and a constant 3’ sequence containing a T7 terminator. Sequences of the 5’ and 3’ constant regions are listed in Supplementary Table 1. Appended sequences were ordered from Twist as Gene Fragments, PCR-amplified, and purified with either Monarch Spin PCR & DNA Cleanup Kit (New England Biolabs T1130) or Sergi Lab magnetic beads (Cat#1040). For the magnetic-bead-based DNA purification, 0.7x of the bead stock solution was added to the PCR product, followed by vigorous mixing with a pipette, incubation at room temperature for 30 minutes to allow complete DNA-to-bead binding, and then by ethanol washes, bead drying and elution as specified in the manufacturer’s protocol. Eluted PCR products were used as the DNA template for in vitro protein synthesis, using PURExpress In Vitro Protein Synthesis Kit (New England Biolabs E6800) with 2 hours of incubation at 37 °C. Reaction products proceeded immediately to the downstream assays within the same day.

### LuxSit luciferase assay

A stock solution of diphenylterazine (DTZ; MedChemExpress #HY-111382) was prepared in N,N-dimethylformamide (DMF) at 10 mM, aliquoted into single-use portions and frozen at -80 °C. After protein synthesis, 50 µL of reaction products was purified with NEBExpress Ni Spin Columns (New England Biolabs S1427) to remove His-tagged transcription and translation machineries from PURExpress, with 30 minutes incubation with the Ni resin at 4 °C on a tube rotator. Purified proteins were stored at 4 °C and used for the luciferase assay within 1 hour. For the luciferase assay, 10 µL of the purified protein was loaded into a well containing 40 µL of 1x PBS in a transparent flat-bottom 96-well plate. Different luciferase variants are separated by at least 1 empty well apart, to minimize signal crosstalk between variants. A 2x assay buffer was prepared by diluting the 10 mM DTZ stock solution into 1x PBS to an intermediate concentration of 50 µM (200-fold dilution). 50 µL of the 2x assay buffer was added to each well, resulting in 100 µL per well and a final DTZ concentration of 25 µM. The 96-well plate was immediately loaded into an Agilent Biotek Cytation 5 reader, with a set temperature of 25 °C. Luminescence was read every minute for 2 hours, with 3 seconds of orbital shaking (5-mm displacement; 307 cycles per minute) prior to each read.

### Uricase assay

Uricase variants were synthesized according to the section “General procedure for synthesizing enzymes for activity assays”, with the modification that the 3’ constant region appended to the uricase sequences additionally encodes a C-terminal fusion of a GGSSG linker followed by a HiBiT tag (sequence available in Supplementary Table 1). A stock solutions of 100 mM sodium 3,5-dichloro-2-hydroxybenzenesulfonate (DCHBS; Millipore Sigma D4645) and 50 mM 4-aminoantipyrine (4-AAP; Millipore Sigma A4382) were prepared in deionized water, aliquoted into single-use portions and frozen at -20 °C. Horseradish peroxidase Type VI (Millipore Sigma P8375) was prepared into a stock solution of 250 units/mL in 100 mM Tris-HCl pH 7.5 + 50% glycerol, aliquoted and stored at -20 °C. Stock solution of 10 mM uric acid (Millipore Sigma U2625) was prepared freshly on the day of experiment, by first dissolving the uric acid powder in 0.1 M NaOH (0.5 mL per 1 mg uric acid), and then adding deionized water until the concentration reached 10 mM. After protein synthesis, uricases were first analyzed with a HiBiT-mediated quantification assay. 1 µL of reaction products was diluted by 30,000-fold in 1x PBS + 0.2 mg/mL bovine serum albumin. 50 µL of the diluted products was loaded on a white, round-bottom 96-well plate. Each well was added with 50 µL of Nano-Glo HiBiT Lytic Reagent (Promega N3030), which contains the HiBiT-complementary LgBiT protein and the luciferin for the reconstituted luciferase. The 96-well plate was immediately loaded into an Agilent Biotek Cytation 5 reader, with a set temperature of 25 °C. The plate underwent 5 minutes of orbital shaking (5-mm displacement; 307 cycles per minute) and 15 minutes of static incubation at 25 °C, after which luminescence was recorded.

After HiBiT-mediated quantification, uricases were analyzed with an activity assay. 2 µL of the protein synthesis reaction products was loaded into a transparent flat-bottom 96-well plate. An assay buffer containing 100 mM Tris-HCl pH 8.6, 4 mM DCHBS, 1 mM 4-AAP, 0.1 mM uric acid, and 0.1 unit/mL horseradish peroxidase was prepared from stock solutions specified in the previous paragraph. Each well was added with 198 µL of the assay buffer, and the 96-well plate was immediately loaded into an Agilent Biotek Cytation 5 reader, with a set temperature of 37 °C. Absorbance at 520 nm was read every minute for 2 hours, with 5 seconds of orbital shaking (5-mm displacement; 307 cycles per minute) prior to each read.

### Computational details of MESSI

#### Set up

MESSI is an inference-time steering method to guide a frozen discrete diffusion protein language model with trained preresentation risk heads.

#### A whole-sequence model of presentation risk

MHC-II presentation is a property of overlapping peptide windows instead of isolated residues: a single substitution rewrites every window that spans it, two substitutions closer together than one window length interact explicitly, and the same residues also encode folding, catalysis and molecular recognition. Deimmunization is therefore constrained search over complete sequences rather than independent deletion of predicted epitopes, which is the regime that earlier sequence- and structure-based optimization methods were built for (*1, 2*).

Let *S* be a target backbone, *a* an HLA-II allele, and *y* = (*y*_1_, …, *y_L_*) e A*^L^* a complete sequence over the twenty amino acids A. Positions are partitioned before design into immutable anchors *C*, with specified residue identities, and an editable set *E* = (1, …, *L*} \ *C*. An immutable complete native reference *y̅* is supplied for the cumulative window-deterioration constraint. For this fixed backbone, allele, reference and constraint specification, we use 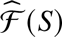 as shorthand for the sequences that preserve the anchors, pass the declared structural gate, and satisfy the window-level constraint defined below.

A risk model, namely the Head, assigns each complete sequence a scalar predicted presentation burden *R_H_* (*y*; *a*), with lower values preferred. In particular, The Head comprises a multiscale dilated convolutional sequence encoder and a trained span scorer. Contextual residue features are projected to 128 dimensions and used to represent each peptide window through its interior, endpoints, flanks and length, together with a single-entry learned allele embedding within each separately trained checkpoint. The encoder and scorer are trained jointly using span supervision, IoU-graded ranking and residue-level supervision. Separate production checkpoints define the reported allele-specific objectives and are frozen before design.

For allele *a*, the corresponding Head scores every contiguous window *w* of 12 to 25 residues and returns a logit *z_w_*_,*a*_ (*y*). Window logits are aggregated by window-count-normalized log-mean-exp to give a per-residue evaluation and a whole-protein score,

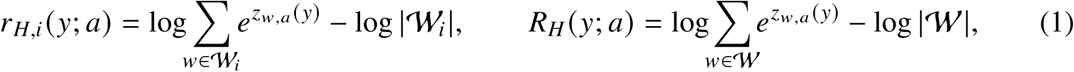

where W is the full window set and W*_i_* = (*w* e W : *i* e *w*} the set of windows containing residue *i*. Subtracting the log-count normalizes for the number of aggregated windows; log-mean-exp retains contributions from multiple windows. The whole-protein score *R_H_* is the sole immune optimization objective in the single-allele formulation, whereas the residue summaries *r_H_*_,*i*_ provide interpretable views of the landscape.

Two properties make this model usable inside a search loop. First, it returns a continuous whole-protein score such that partial progress is visible: a redesign that weakens an epitope without abolishing it registers, whereas a threshold-crossing count does not. Second, it is much faster than traditional evaluation methods: a whole-protein call takes about 0.006 s for a 50-residue protein and 0.11 s for a 1500-residue protein on a single GPU, against roughly 4 s and 150 s respectively for NetMHCIIpan-4.3 on the same sequences and hardware. The per-call speedup therefore rises from about 700-fold at 50 residues to about 1500-fold for proteins longer than 500 residues, the regime that covers most enzyme therapeutics. Because a single MESSI design issues several hundred whole-protein risk evaluations, this speedup ratio is an enabling condition.

#### Structure-conditioned generator

Sequence proposals come from a frozen structure-conditioned discrete diffusion protein language model: a 650-million-parameter masked diffusion language model with a geometric vector perceptron encoder that reads the target backbone once per design and conditions every decoding step (*3*). Generation begins with the editable set fully masked and runs for *T* = 100 denoising steps. At step *t* the denoiser emits logits at all positions; each still-masked position is independently unmasked with a schedule-determined probability, and the identity at each newly unmasked position is drawn from the temperature-scaled categorical distribution at temperature 1.0 to maintain generation diversity. Anchor identities are re-imposed after every step. The same frozen generator, schedule and temperature are used by MESSI and its feedback-disabled boundary. Background global remasking is disabled during ordinary completion and propagation; resolved editable identities can be reopened only by the explicit feedback intervention.

### Design objective

The single-allele design problem is therefore

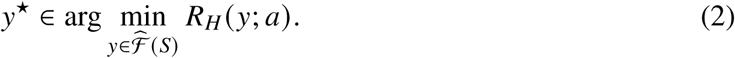

Taking computation resources into account, we report the achievable risk as a resource-indexed frontier,

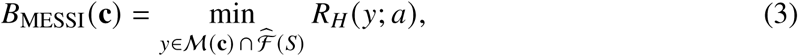

where M(**c**) is the set of complete sequences evaluated under a declared resource vector **c** whose coordinates are denoiser evaluations, Head calls, structure evaluations, GPU-hours and wall time.

For joint two-allele design, two separately trained Heads score the same exact sequence. Frozen affine calibrations define *u_a_* (*y*) = [*R_a_* (*y*) − *b_a_*]/*s_a_*, with *s_a_* > 0, and the objective is

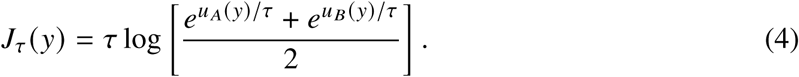

The calibration constants and *τ* are specified before search. The same joint objective is used for donor, incumbent and archive comparisons and for local reverts: the identical reverted sequence is scored by both Heads, and its contribution is *J_τ_* (*y*^(−^*^i^*^)^) − *J_τ_* (*y*). Per-allele window constraints remain explicit. A lower joint objective does not imply that both allele scores decrease; paired allele-specific outcomes are reported separately.

### Partial sequences as search states

Adding a mask symbol ⊥, a partial sequence 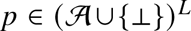 denotes the set of complete sequences that agree with it,

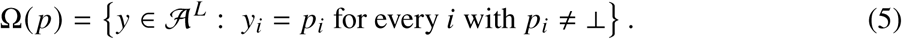

Ordered by inclusion, these cylinders form a partially ordered set. Fixing an identity contracts the set of syntactically compatible completions, returning a position to ⊥ expands it, and leaving a position unchanged preserves it. This set-level description is distinct from the probabilities assigned by the generator: compatibility alone does not establish positive sampling probability.

A token array alone does not determine a legal continuation. Two states with identical visible residues but different generator times, or different histories of how those residues came to be fixed, admit different next steps. The executable state is therefore

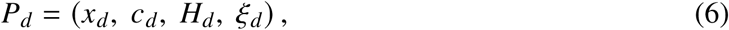

with *S*, *C* and *E* fixed for the run. Here *x_d_* is the partial sequence, *c_d_* e (0, …, *T*} is the generator time at which it was captured, *ξ_d_* is the random state that permits exact replay, and *H_d_* records, for each position, when it was resolved, whether its identity was produced by the generator or written by feedback, and whether it has yet been scored in its current context. *H_d_* is not supplied as a learned input to the neural denoiser, but consumed by projection legality, active-score bookkeeping and subsequent support selection, and retains the origin information needed to audit the intervention.

#### Algorithmic details

MESSI advances one active trajectory through at most *D* committed feedback cycles. Each cycle (i) looks ahead: generates multiple complete sequences; (ii) selects: evaluate these continuations with the Head and filters out the single candidate with lowest risk; (iii) attributes: evaluate residue-level risk of the donor; and (iv) projects: partially merge the donor into the current state using residue-level adoption or reopening rules.

#### Look ahead

At the capture time *c_d_*, MESSI draws *K_d_* complete continuations from the same unfinished state,

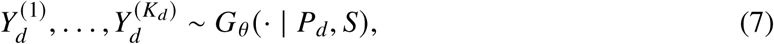

where *G_θ_* runs the frozen denoiser from *c_d_* to completion. The forks use disjoint, replayable seeds and share every residue the source has already committed, so they differ only in the tail the generator has not yet written.

#### Select

Endpoints are scored and gated. Let L*_d_* denote the complete evaluated endpoint pool, in-cluding its admission verdicts, and 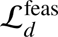 denote the subset passing every declared gate on measured evidence, including structural evaluation. Repeated sequences are collapsed to a single representa-tive before selection. When this subset is nonempty, the donor is the lexicographic minimum

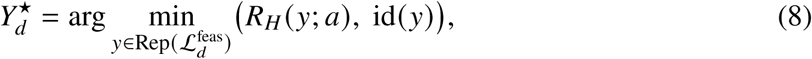

where the endpoint identifier breaks ties deterministically. Every evaluated endpoint enters the archive M*_d_*, while only admitted endpoints may direct the search.

At *d* = 0 the best gated endpoint becomes the lineage incumbent *I*_1_. For *d* ≥ 1, feedback proceeds only if the donor beats the incumbent by a fixed margin,

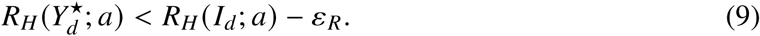

The margin *ε_R_* is fixed before search at twice the maximum absolute drift observed when the frozen Head repeatedly scores identical sequences in a calibration panel. The local-contribution threshold is calibrated from the same repeat-scoring procedure. When no donor clears the margin, the lineage records a typed stall and stops further feedback, while the archive is unaffected.

#### Attribute

To identify residue-level contribution to the improvement, MESSI reverts one donor position *i* that differs from the reference *J_d_* at a time to *J_d_*_,*i*_, leaving the rest of the donor intact:

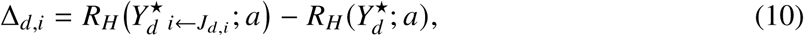

with *J*_0_ the native sequence and *J_d_* = *I_d_* for *d* ≥ 1. A position is eligible only if it is editable, unresolved in the live source, different between donor and reference, and covered by an aligned window that improves in the donor. A value of Δ*_d_*_,*i*_ above the frozen repeatability threshold means that this reversion worsens the Head score in the exact donor context. As a prespecified intervention-size constraint, the number of newly adopted identities is capped at

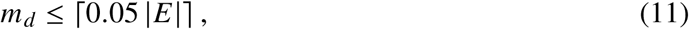

For positions already resolved in the source, compare residue-level risk

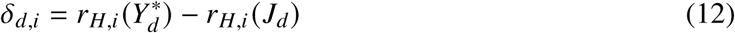

where a positive *δ_d_*_,*i*_ suggests the residue *i* leads to higher risk in the donor context and thus should be regenerated.

#### Project

The projection consists of two parts: adopting some identities by inheriting residues with large Δ*_d_*_,*i*_ in the donor, and remasking (or reopening) some resolved positions with large *δ_d_*_,*i*_. To enable more flexibility in renewing high-risk positions according to the new contexts, after adopting *m_d_* positions from the donor, the sequence re-entries at an earlier time *r_d_* < *c_d_*. If the source has *u*_src_ unresolved editable positions and the target remaining mask positions is *u*_target_, adopting *m_d_* identities requires exactly

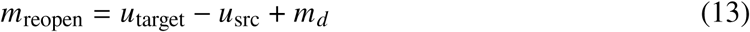

previously resolved editable positions to be reopened to ⊥. Eligible positions are ranked determin-istically by the largest positive increase in covering window logits relative to the immutable native reference, namely *δ_d_*_,*i*_.

The outcome is a partition of the editable positions into four labeled classes,

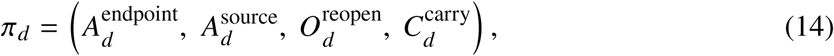

separating identities written from the donor, identities that the source committed at or after *r_d_* and must therefore be re-asserted explicitly rather than assumed, positions returned to ⊥, and positions whose value at *r_d_* is already what the trajectory naturally held. Anchors never enter the partition.

Under the frozen no-background-remask substrate, the same denoiser propagates the state from timestep *r_d_* to *c_d_*_+1_, from which the next complete continuation pool is generated, and the entire procedure above repeats until *T*. Concretely, the projection and the resumed transition are

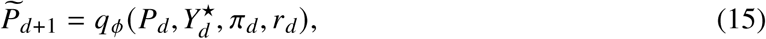

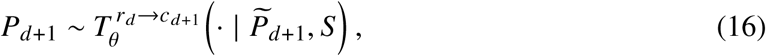

with *r_d_* < *c_d_* and *c_d_*_+1_ ≥ *c_d_*; the primary progressive schedule uses *c_d_*_+1_ > *c_d_*. By default, we set *c_d_* − *r_d_* = 10.

## Algorithm 1 1 Trajectory-coupled endpoint feedback in MESSI

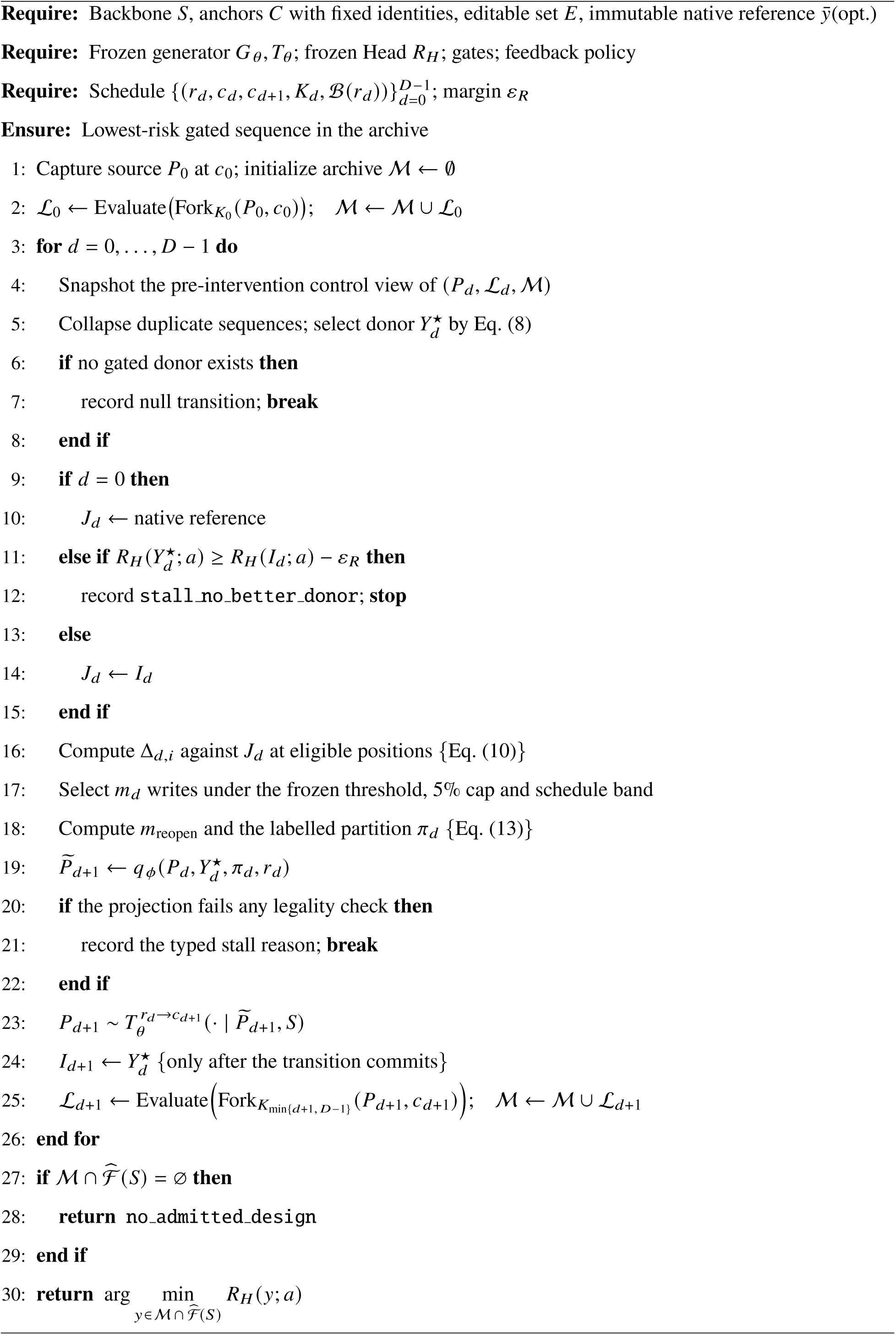

## References and Notes

1. L. Gandhi, D. Rodríguez-Abreu, S. Gadgeel, E. Esteban, E. Felip, F. De Angelis, M. Domine, P. Clingan, M. J. Hochmair, S. F. Powell, S. Y.-S. Cheng, H. G. Bischoff, N. Peled, F. Grossi, R. R. Jennens, M. Reck, R. Hui, E. B. Garon, M. Boyer, B. Rubio-Viqueira, S. Novello, T. Kurata, J. E. Gray, J. Vida, Z. Wei, J. Yang, H. Raftopoulos, M. C. Pietanza, M. C. Garassino, KEYNOTE-189 Investigators, Pembrolizumab plus chemotherapy in metastatic non-small-cell lung cancer. N. Engl. J. Med. 378, 2078–2092 (2018).

2. M. Kothari, A. Wanjari, S. Acharya, V. Karwa, R. Chavhan, S. Kumar, A. Kadu, R. Patil, A comprehensive review of monoclonal antibodies in modern medicine: Tracing the evolution of a revolutionary therapeutic approach. Cureus 16, e61983 (2024).

3. P. J. Carter, V. Quarmby, Immunogenicity risk assessment and mitigation for engineered antibody and protein therapeutics. Nat. Rev. Drug Discov. 23, 898–913 (2024).

4. P. A. Roche, K. Furuta, The ins and outs of MHC class II-mediated antigen processing and presentation. Nat. Rev. Immunol. 15, 203–216 (2015).

5. A. Vaisman-Mentesh, M. Gutierrez-Gonzalez, B. J. DeKosky, Y. Wine, The molecular mechanisms that underlie the immune biology of anti-drug antibody formation following treatment with monoclonal antibodies. Front. Immunol. 11, 1951 (2020).

6. M. P. Baker, H. M. Reynolds, B. Lumicisi, C. J. Bryson, Immunogenicity of protein therapeutics: The key causes, consequences and challenges. Self Nonself 1, 314–322 (2010).

7. J. Link, M. Lundkvist Ryner, K. Fink, C. Hermanrud, I. Lima, B. Brynedal, I. Kockum, J. Hillert, A. Fogdell-Hahn, Human leukocyte antigen genes and interferon beta preparations influence risk of developing neutralizing anti-drug antibodies in multiple sclerosis. PLoS One 9, e90479 (2014).

8. S. Malucchi, A. Sala, F. Gilli, R. Bottero, A. Di Sapio, M. Capobianco, A. Bertolotto, Neutralizing antibodies reduce the efficacy of betaIFN during treatment of multiple sclerosis. Neurology 62, 2031–2037 (2004).

9. C. Witmer, G. Young, Factor VIII inhibitors in hemophilia A: rationale and latest evidence. Ther. Adv. Hematol. 4, 59–72 (2013).

10. I. M. van der Sluis, L. M. Vrooman, R. Pieters, A. Baruchel, G. Escherich, N. Goulden, V. Mondelaers, J. Sanchez de Toledo, C. Rizzari, L. B. Silverman, J. A. Whitlock, Consensus expert recommendations for identification and management of asparaginase hypersensitivity and silent inactivation. Haematologica 101, 279–285 (2016).

11. J. R. Cantor, T. H. Yoo, A. Dixit, B. L. Iverson, T. G. Forsthuber, G. Georgiou, Therapeutic enzyme deimmunization by combinatorial T-cell epitope removal using neutral drift. Proc. Natl. Acad. Sci. U. S. A. 108, 1272–1277 (2011).

12. P. E. Lipsky, L. H. Calabrese, A. Kavanaugh, J. S. Sundy, D. Wright, M. Wolfson, M. A. Becker, Pegloticase immunogenicity: the relationship between efficacy and antibody development in patients treated for refractory chronic gout. Arthritis Res. Ther. 16, R60 (2014).

13. K. C. Allen, A. H. Champlain, J. A. Cotliar, S. M. Belknap, D. P. West, J. Mehta, S. M. Trifilio, Risk of anaphylaxis with repeated courses of rasburicase: a Research on Adverse Drug Events and Reports (RADAR) project. Drug Saf. 38, 183–187 (2015).

14. M.-S. E. D. Schulze, K. W. Wucherpfennig, The mechanism of HLA-DM induced peptide exchange in the MHC class II antigen presentation pathway. Curr. Opin. Immunol. 24, 105–111 (2012).

15. V. Jawa, L. P. Cousens, M. Awwad, E. Wakshull, H. Kropshofer, A. S. De Groot, T-cell dependent immunogenicity of protein therapeutics: Preclinical assessment and mitigation. Clin. Immunol. 149, 534–555 (2013).

16. V. Jawa, F. Terry, J. Gokemeijer, S. Mitra-Kaushik, B. J. Roberts, S. Tourdot, A. S. De Groot, T-cell dependent immunogenicity of protein therapeutics pre-clinical assessment and mitigation-updated consensus and review 2020. Front. Immunol. 11, 1301 (2020).

17. J. H. Brown, T. S. Jardetzky, J. C. Gorga, L. J. Stern, R. G. Urban, J. L. Strominger, D. C. Wiley, Three-dimensional structure of the human class II histocompatibility antigen HLA-DR1. Nature 364, 33–39 (1993).

18. L. J. Stern, J. H. Brown, T. S. Jardetzky, J. C. Gorga, R. G. Urban, J. L. Strominger, D. C. Wiley, Crystal structure of the human class II MHC protein HLA-DR1 complexed with an influenza virus peptide. Nature 368, 215–221 (1994).

19. B. Magyari, A. T. Fülöp, D. Kókai, F. Tóth, G. M. Balogh, S. Andreu-Sánchez, G. Innocenti, R. K. Weersma, J. Fu, C. Pál, A. Zhernakova, T. Vogl, M. Manczinger, HLA-II peptide-binding diversity shapes humoral immune responses and susceptibility to infections. Nat. Commun. 17, 9422 (2026).

20. R. M. Chicz, R. G. Urban, J. C. Gorga, D. A. Vignali, W. S. Lane, J. L. Strominger, Specificity and promiscuity among naturally processed peptides bound to HLA-DR alleles. J. Exp. Med. 178, 27–47 (1993).

21. J. Racle, P. Guillaume, J. Schmidt, J. Michaux, A. Larabi, K. Lau, M. A. S. Perez, G. Croce, R. Genolet, G. Coukos, V. Zoete, F. Pojer, M. Bassani-Sternberg, A. Harari, D. Gfeller, Machine learning predictions of MHC-II specificities reveal alternative binding mode of class II epitopes. Immunity 56, 1359–1375.e13 (2023).

22. C. G. Rappazzo, B. D. Huisman, M. E. Birnbaum, Repertoire-scale determination of class II MHC peptide binding via yeast display improves antigen prediction. Nat. Commun. 11, 4414 (2020).

23. F. A. Harding, M. M. Stickler, J. Razo, R. B. DuBridge, The immunogenicity of humanized and fully human antibodies: residual immunogenicity resides in the CDR regions: Residual immunogenicity resides in the CDR regions. MAbs 2, 256–265 (2010).

24. H. Stark, F. Faltings, M. Choi, Y. Xie, E. Hur, T. O’Donnell, A. Bushuiev, T. Uçar, S. Passaro, W. Mao, M. Reveiz, R. Bushuiev, T. Pluskal, J. Sivic, K. Kreis, A. Vahdat, S. Ray, J. T. Goldstein, A. Savinov, J. A. Hambalek, A. Gupta, D. A. Taquiri-Diaz, Y. Zhang, A. K. Hatstat, A. Arada, N. H. Kim, E. Tackie-Yarboi, D. Boselli, L. Schnaider, C. C. Liu, G.-W. Li, D. Hnisz, D. M. Sabatini, W. F. DeGrado, J. Wohlwend, G. Corso, R. Barzilay, T. Jaakkola, BoltzGen: Toward universal binder design, bioRxivorg (2025). 10.1101/2025.11.20.689494.

25. Y. Zhang, C. Gong, J. Sun, J. Guan, M. Ren, S. Xue, H. Zhang, W. Ma, Z. Liu, X. Chen, W. Xiao, Protenix-v2: Broadening the reach of structure prediction and biomolecular design, bioRxiv (2026)p. 2026.04.10.717613.

26. L. S. Mille-Fragoso, C. L. Driscoll, J. N. Wang, H. Dai, T. Widatalla, J. L. Zhang, X. Zhang, B. Rao, L. Feng, B. L. Hie, X. J. Gao, Efficient generation of epitope-targeted antibodies with Germinal. Nat. Biotechnol., doi: 10.1038/s41587-026-03187-0 (2026).

27. M. Pacesa, L. Nickel, C. Schellhaas, J. Schmidt, E. Pyatova, L. Kissling, P. Barendse, J. Choudhury, S. Kapoor, A. Alcaraz-Serna, Y. Cho, K. H. Ghamary, L. Vinué, B. J. Yachnin, A. M. Wollacott, S. Buckley, A. H. Westphal, S. Lindhoud, S. Georgeon, C. A. Goverde, G. N. Hatzopoulos, P. Gönczy, Y. D. Muller, G. Schwank, D. C. Swarts, A. J. Vecchio, B. L. Schneider, S. Ovchinnikov, B. E. Correia, One-shot design of functional protein binders with BindCraft. Nature 646, 483–492 (2025).

28. J. L. Watson, D. Juergens, N. R. Bennett, B. L. Trippe, J. Yim, H. E. Eisenach, W. Ahern, A. J. Borst, R. J. Ragotte, L. F. Milles, B. I. M. Wicky, N. Hanikel, S. J. Pellock, A. Courbet, W. Sheffler, J. Wang, P. Venkatesh, I. Sappington, S. V. Torres, A. Lauko, V. De Bortoli, E. Mathieu, S. Ovchinnikov, R. Barzilay, T. S. Jaakkola, F. DiMaio, M. Baek, D. Baker, De novo design of protein structure and function with RFdiffusion. Nature 620, 1089–1100 (2023).

29. N. R. Bennett, J. L. Watson, R. J. Ragotte, A. J. Borst, D. L. See, C. Weidle, R. Biswas, Y. Yu, E. L. Shrock, R. Ault, P. J. Y. Leung, B. Huang, I. Goreshnik, J. Tam, K. D. Carr, B. Singer, C. Criswell, B. I. M. Wicky, D. Vafeados, M. Garcia Sanchez, H. M. Kim, S. Vázquez Torres, S. Chan, S. M. Sun, T. T. Spear, Y. Sun, K. O’Reilly, J. M. Maris, N. G. Sgourakis, R. A. Melnyk, C. C. Liu, D. Baker, Atomically accurate de novo design of antibodies with RFdiffusion. Nature 649, 183–193 (2026).

30. S. Candido, T. Hayes, A. Derry, R. Rao, Z. Lin, R. Verkuil, B. Z. Wu, J. S. Lee, E. S. Bruguera, J. A. Keval, M. Kopylov, J. E. Pak, W. Wu, N. Thomas, S. Mataraso, A. Hsu, A. C. Trotman-Grant, K. Fatras, A. dos Santos Costa, R. Badkundri, H. Akın, D. Oktay, J. Deaton, E. Montabana, H. Sitwala, Y. Yu, M. Wiggert, D. A. Carlin, A. W. Goering, T. Blazejewski, M. Sandora, M. Hla, T. Z. Jia, L. H. Kloker, N. J. Sofroniew, M. Uehara, J. Pannu, S. Bachas, D. S. Liu, T. Sercu, A. Rives, Language modeling materializes a world model of protein biology, bioRxiv (2026)p. 2026.06.03.729735.

31. W. J. Thrift, J. Perera, S. Cohen, N. W. Lounsbury, H. R. Gurung, C. M. Rose, J. Chen, S. Jhunjhunwala, K. Liu, Graph-pMHC: graph neural network approach to MHC class II peptide presentation and antibody immunogenicity. Brief. Bioinform. 25, bbae123 (2024).

32. M. Haghighatlari, N. Marze, R. Seward, A. Ciarla, R. Hindin, J. Calderini, B. Keenan, S. Dhule, S. Hall-Swan, T. P. Hickling, E. Bennett, B. Rai, S. Tourdot, HLAIIPred: cross-attention mechanism for modeling the interaction of HLA class II molecules with peptides. *Commun*. Biol. 8, 1133 (2025).

33. J. B. Nilsson, S. Kaabinejadian, H. Yari, M. G. D. Kester, P. van Balen, W. H. Hildebrand, M. Nielsen, Accurate prediction of HLA class II antigen presentation across all loci using tailored data acquisition and refined machine learning. Sci. Adv. 9, eadj6367 (2023).

34. E. Wolfsberg, J.-S. Paul, J. Tycko, B. Chen, M. C. Bassik, L. Bintu, A. A. Alizadeh, X. J. Gao, Machine-guided dual-objective protein engineering for deimmunization and therapeutic functions. Cell Syst. 16, 101299 (2025).

35. B. Schubert, C. Schärfe, P. Dönnes, T. Hopf, D. Marks, O. Kohlbacher, Population-specific design of de-immunized protein biotherapeutics. PLoS Comput. Biol. 14, e1005983 (2018).

36. C. King, E. N. Garza, R. Mazor, J. L. Linehan, I. Pastan, M. Pepper, D. Baker, Removing T-cell epitopes with computational protein design. Proc. Natl. Acad. Sci. U. S. A. 111, 8577–8582 (2014).

37. A. S. Parker, Y. Choi, K. E. Griswold, C. Bailey-Kellogg, Structure-guided deimmunization of therapeutic proteins. J. Comput. Biol. 20, 152–165 (2013).

38. R. S. Salvat, Y. Choi, A. Bishop, C. Bailey-Kellogg, K. E. Griswold, Protein deimmunization via structure-based design enables efficient epitope deletion at high mutational loads: Efficient Structure-Based Protein Deimmunization. Biotechnol. Bioeng. 112, 1306–1318 (2015).

39. R. S. Salvat, D. Verma, A. S. Parker, J. R. Kirsch, S. A. Brooks, C. Bailey-Kellogg, K. E. Griswold, Computationally optimized deimmunization libraries yield highly mutated enzymes with low immunogenicity and enhanced activity. Proc. Natl. Acad. Sci. U. S. A. 114, E5085–E5093 (2017).

40. Y. Choi, D. Verma, K. E. Griswold, C. Bailey-Kellogg, EpiSweep: Computationally driven reengineering of therapeutic proteins to reduce immunogenicity while maintaining function. Methods Mol. Biol. 1529, 375–398 (2017).

41. Z. Lin, H. Akin, R. Rao, B. Hie, Z. Zhu, W. Lu, N. Smetanin, R. Verkuil, O. Kabeli, Y. Shmueli, A. Dos Santos Costa, M. Fazel-Zarandi, T. Sercu, S. Candido, A. Rives, Evolutionary-scale prediction of atomic-level protein structure with a language model. Science 379, 1123–1130 (2023).

42. T. Hayes, R. Rao, H. Akin, N. J. Sofroniew, D. Oktay, Z. Lin, R. Verkuil, V. Q. Tran, J. Deaton, M. Wiggert, R. Badkundri, I. Shafkat, J. Gong, A. Derry, R. S. Molina, N. Thomas, Y. A. Khan, C. Mishra, C. Kim, L. J. Bartie, M. Nemeth, P. D. Hsu, T. Sercu, S. Candido, A. Rives, Simulating 500 million years of evolution with a language model. Science 387, 850–858 (2025).

43. J. Jumper, R. Evans, A. Pritzel, T. Green, M. Figurnov, O. Ronneberger, K. Tunyasuvunakool, R. Bates, A. Žídek, A. Potapenko, A. Bridgland, C. Meyer, S. A. A. Kohl, A. J. Ballard, A. Cowie, B. Romera-Paredes, S. Nikolov, R. Jain, J. Adler, T. Back, S. Petersen, D. Reiman, E. Clancy, M. Zielinski, M. Steinegger, M. Pacholska, T. Berghammer, S. Bodenstein, D. Silver, O. Vinyals, A. W. Senior, K. Kavukcuoglu, P. Kohli, D. Hassabis, Highly accurate protein structure prediction with AlphaFold. Nature 596, 583–589 (2021).

44. J. Abramson, J. Adler, J. Dunger, R. Evans, T. Green, A. Pritzel, O. Ronneberger, L. Willmore, A. J. Ballard, J. Bambrick, S. W. Bodenstein, D. A. Evans, C.-C. Hung, M. O’Neill, D. Reiman, K. Tunyasuvunakool, Z. Wu, A. Žemgulytė, E. Arvaniti, C. Beattie, O. Bertolli, A. Bridgland, A. Cherepanov, M. Congreve, A. I. Cowen-Rivers, A. Cowie, M. Figurnov, F. B. Fuchs, H. Gladman, R. Jain, Y. A. Khan, C. M. R. Low, K. Perlin, A. Potapenko, P. Savy, S. Singh, A. Stecula, A. Thillaisundaram, C. Tong, S. Yakneen, E. D. Zhong, M. Zielinski, A. Žídek, V. Bapst, P. Kohli, M. Jaderberg, D. Hassabis, J. M. Jumper, Accurate structure prediction of biomolecular interactions with AlphaFold 3. Nature 630, 493–500 (2024).

45. V. R. Shanker, T. U. J. Bruun, B. L. Hie, P. S. Kim, Unsupervised evolution of protein and antibody complexes with a structure-informed language model. Science 385, 46–53 (2024).

46. P. E. Cavanagh, A. G. Xue, S. A. Dai, A. Qiang, T. Matsui, A. Y. Ting, Computational design of conformation-biasing mutations to alter protein functions. Science 391, eadv7953 (2026).

47. P. Skopintsev, I. Esain-Garcia, E. C. DeTurk, P. H. Yoon, Z. Zhou, T. Weiss, M. Kamalu, A. Chamraj, K. J. Loi, C. J. Langeberg, R. S. Boger, H. Nisonoff, H. M. Karp, L.-X. Chen, H. Shi, K. Vohra, J. F. Banfield, J. H. D. Cate, S. E. Jacobsen, J. A. Doudna, Structure and evolution-guided design of minimal RNA-guided nucleases. Science 393, 313–318 (2026).

48. X. Wang, Z. Zheng, F. Ye, D. Xue, S. Huang, Q. Gu, Diffusion language models are versatile protein learners, arXiv [cs.LG] (2024). 10.48550/arXiv.2402.18567.

49. X. Wang, Z. Zheng, F. Ye, D. Xue, S. Huang, Q. Gu, DPLM-2: A Multimodal Diffusion Protein Language Model, arXiv [cs.LG] (2024). 10.48550/arXiv.2410.13782.

50. J. Dauparas, I. Anishchenko, N. Bennett, H. Bai, R. J. Ragotte, L. F. Milles, B. I. M. Wicky, A. Courbet, R. J. de Haas, N. Bethel, P. J. Y. Leung, T. F. Huddy, S. Pellock, D. Tischer, F. Chan, B. Koepnick, H. Nguyen, A. Kang, B. Sankaran, A. K. Bera, N. P. King, D. Baker, Robust deep learning-based protein sequence design using ProteinMPNN. Science 378, 49–56 (2022).

51. C. Hsu, R. Verkuil, J. Liu, Z. Lin, B. Hie, T. Sercu, A. Lerer, A. Rives, Learning inverse folding from millions of predicted structures, bioRxiv (2022)p. 2022.04.10.487779.

52. P. Notin, A. W. Kollasch, D. Ritter, L. van Niekerk, S. Paul, H. Spinner, N. Rollins, A. Shaw, R. Weitzman, J. Frazer, M. Dias, D. Franceschi, R. Orenbuch, Y. Gal, D. S. Marks, ProteinGym: Large-scale benchmarks for protein design and fitness prediction, bioRxivorg (2023). 10.1101/2023.12.07.570727.

53. J. Rector-Brooks, T. Lambert, M. Skreta, D. Roth, Y. Long, Z.-Q. Li, X. Zhang, M. Cretu, F.-Z. Li, T. Ganapathy, E. Jin, A. J. Bose, J. Yang, K. Neklyudov, Y. Bengio, A. Tong, F. H. Arnold, C.-H. Liu, General multimodal protein design enables DNA-encoding of chemistry, arXiv [cs.LG] (2026). 10.48550/arXiv.2604.05181.

54. H.-C. Gasser, D. A. Oyarzún, A. Rajan, J. A. Alfaro, Guiding a language-model based protein design method towards MHC Class-I immune-visibility targets in vaccines and therapeutics. Immunoinformatics (Amst.) 14, 100035 (2024).

55. H.-C. Gasser, A. Rajan, J. A. Alfaro, A novel decoding strategy for ProteinMPNN to design with less visibility to cytotoxic T-lymphocytes. Comput. Struct. Biotechnol. J. 27, 3693–3703 (2025).

56. S. Tang, Y. Zhu, M. Tao, P. Chatterjee, TR2-D2: Tree Search Guided TRajectory-Aware Fine-tuning for Discrete Diffusion, ArXiv (2025). 10.48550/arXiv.2509.25171.

57. S. Tang, Y. Zhu, M. Tao, P. Chatterjee, A2D2: Fine-Tuning Any-Length Discrete Diffusion for Adaptive Decoding, arXiv [cs.LG] (2026). 10.48550/arXiv.2606.13565.

58. S. Alamdari, N. Thakkar, R. van den Berg, N. Tenenholtz, R. Strome, A. M. Moses, A. X. Lu, N. Fusi, A. P. Amini, K. K. Yang, Protein generation with evolutionary diffusion: sequence is all you need, eLife (2026). 10.7554/elife.112029.1.

59. K. E. Wu, K. K. Yang, R. van den Berg, S. Alamdari, J. Y. Zou, A. X. Lu, A. P. Amini, Protein structure generation via folding diffusion. Nat. Commun. 15, 1059 (2024).

60. W. S. Lu, X. Zhang, L. Mille-Fragoso, H. Dai, X. Gao, W. H. Wong, ProVADA: Generation of subcellular protein variants via ensemble-guided test-time steering, bioRxiv (2025). 10.1101/2025.07.11.664238.

61. J. Austin, D. D. Johnson, J. Ho, D. Tarlow, R. van den Berg, Structured denoising diffusion models in discrete state-spaces, arXiv [cs.LG] (2021). 10.48550/arXiv.2107.03006.

62. N. Gruver, S. Stanton, N. C. Frey, T. G. J. Rudner, I. Hotzel, J. Lafrance-Vanasse, A. Rajpal, K. Cho, A. G. Wilson, Protein design with guided discrete diffusion, arXiv [cs.LG] (2023). 10.48550/arXiv.2305.20009.

63. H. Nisonoff, J. Xiong, S. Allenspach, J. Listgarten, Unlocking guidance for discrete state-space diffusion and flow models, arXiv [cs.LG] (2024). 10.48550/arXiv.2406.01572.

64. H. Stark, B. Jing, C. Wang, G. Corso, B. Berger, R. Barzilay, T. Jaakkola, Dirichlet flow matching with applications to DNA sequence design. ArXiv (2024).

65. S. Tang, Y. Zhang, P. Chatterjee, PepTune: De Novo generation of therapeutic peptides with multi-objective-guided discrete diffusion, arXiv [q-bio.BM] (2024). 10.48550/arXiv.2412.17780.

66. X. Li, Y. Zhao, C. Wang, G. Scalia, G. Eraslan, S. Nair, T. Biancalani, S. Ji, A. Regev, S. Levine, M. Uehara, Derivative-free guidance in continuous and discrete diffusion models with soft value-based decoding, arXiv [cs.LG] (2024). 10.48550/arXiv.2408.08252.

67. G. Wang, Y. Schiff, S. S. Sahoo, V. Kuleshov, Remasking discrete diffusion models with inference-time scaling, arXiv [cs.LG] (2025). 10.48550/arXiv.2503.00307.

68. M. Uehara, X. Su, Y. Zhao, X. Li, A. Regev, S. Ji, S. Levine, T. Biancalani, Reward-guided iterative refinement in diffusion models at test-time with applications to protein and DNA design, arXiv [cs.LG] (2025). 10.48550/arXiv.2502.14944.

69. C. Wang, M. Uehara, Y. He, A. Wang, T. Biancalani, A. Lal, T. Jaakkola, S. Levine, H. Wang, A. Regev, Fine-tuning discrete diffusion models via reward optimization with applications to DNA and protein design, arXiv [cs.LG] (2024). 10.48550/arXiv.2410.13643.

70. P. Phunyaphibarn, M. Sung, Reward-guided discrete diffusion via clean-Sample Markov Chain for molecule and biological sequence design, arXiv [cs.LG] (2026). 10.48550/arXiv.2602.09424.

71. P. Holderrieth, D. Chen, L. Eyring, I. Shah, G. Anantharaman, Y. He, Z. Akata, T. Jaakkola, N. M. Boffi, M. Simchowitz, Diamond Maps: Efficient reward alignment via stochastic flow maps, arXiv [cs.LG] (2026). 10.48550/arXiv.2602.05993.

72. K. Jeon, T.-D. Vuong, M. Tao, VGB for Masked Diffusion Model: Efficient test-time scaling for reward satisfaction and sample editing, arXiv [cs.LG] (2026). 10.48550/arXiv.2606.28301.

73. A. Wachi, T. Tanabe, Y. Akimoto, Safety hacking in constrained Best-of-N inference-time scaling, arXiv [cs.LG] (2026). 10.48550/arXiv.2608.22915.

74. W. C. Hallows, K. Skvorak, N. Agard, N. Kruse, X. Zhang, Y. Zhu, R. C. Botham, C. Chng, C. Shukla, J. Lao, M. Miller, A. Sero, J. Viduya, M. H. A. Ismaili, K. McCluskie, R. Schiffmann, A. P. Silverman, J.-S. Shen, G. W. Huisman, Optimizing human α-galactosidase for treatment of Fabry disease. Sci. Rep. 13, 4748 (2023).

75. R. Vita, N. Blazeska, D. Marrama, IEDB Curation Team Members, S. Duesing, J. Bennett, J. Greenbaum, M. De Almeida Mendes, J. Mahita, D. K. Wheeler, J. R. Cantrell, J. A. Overton, D. A. Natale, A. Sette, B. Peters, The Immune Epitope Database (IEDB): 2024 update. Nucleic Acids Res. 53, D436–D443 (2025).

76. A. H.-W. Yeh, C. Norn, Y. Kipnis, D. Tischer, S. J. Pellock, D. Evans, P. Ma, G. R. Lee, J. Z. Zhang, I. Anishchenko, B. Coventry, L. Cao, J. Dauparas, S. Halabiya, M. DeWitt, L. Carter, K. N. Houk, D. Baker, De novoj design of luciferases using deep learning. Nature 614, 774–780 (2023).

77. K. Henzler-Wildman, D. Kern, Dynamic personalities of proteins. Nature 450, 964–972 (2007).

78. K. Jiang, Z. Yan, M. Di Bernardo, S. R. Sgrizzi, L. Villiger, A. Kayabolen, B. J. Kim, J. K. Carscadden, M. Hiraizumi, H. Nishimasu, J. S. Gootenberg, O. O. Abudayyeh, Rapid in silico directed evolution by a protein language model with EVOLVEpro. Science 387, eadr6006 (2025).

79. T. A. Hopf, J. B. Ingraham, F. J. Poelwijk, C. P. I. Schärfe, M. Springer, C. Sander, D. S. Marks, Mutation effects predicted from sequence co-variation. Nat. Biotechnol. 35, 128–135 (2017).

80. Chai Discovery Team, J. Boitreaud, J. Dent, D. Geisz, M. McPartlon, J. Meier, Z. Qiao, A. Rogozhnikov, N. Rollins, P. Wollenhaupt, K. Wu, Zero-shot antibody design in a 24-well plate, bioRxiv (2025)p. 2025.07.05.663018.

81. D. Bertoni, M. Tsenkov, P. Magana, S. Nair, I. Pidruchna, M. Querino Lima Afonso, A. Midlik, U. Paramval, D. Lawal, A. Tanweer, M. Last, R. Patel, A. Laydon, D. Lasecki, N. Dietrich, H. Tomlinson, A. Žídek, T. Green, O. Kovalevskiy, A. Lau, S. Kandathil, N. Bordin, I. Sillitoe, M. Mirdita, D. Jones, C. Orengo, M. Steinegger, J. R. Fleming, S. Velankar, AlphaFold Protein Structure Database 2025: a redesigned interface and updated structural coverage. Nucleic Acids Res. 54, D358–D362 (2026).

82. F. McCush, E. Wang, C. Yunis, P. Schwartz, D. Baltrukonis, Anti-drug antibody magnitude and clinical relevance using signal to noise (S/N): Bococizumab case study. AAPS J. 25, 85 (2023).

83. E. L. Kneepkens, J. C.-C. Wei, M. T. Nurmohamed, K.-J. Yeo, C. Y. Chen, I. E. van der Horst-Bruinsma, D. van der Kleij, T. Rispens, G. Wolbink, C. L. M. Krieckaert, Immunogenicity, adalimumab levels and clinical response in ankylosing spondylitis patients during 24 weeks of follow-up. Ann. Rheum. Dis. 74, 396–401 (2015).

84. Y. Han, D. Liu, L. Li, PD-1/PD-L1 pathway: current researches in cancer. Am. J. Cancer Res. 10, 727–742 (2020).

85. C. A. Goverde, M. Pacesa, N. Goldbach, L. J. Dornfeld, P. E. M. Balbi, S. Georgeon, S. Rosset, S. Kapoor, J. Choudhury, J. Dauparas, C. Schellhaas, S. Kozlov, D. Baker, S. Ovchinnikov, A. J. Vecchio, B. E. Correia, Computational design of soluble and functional membrane protein analogues. Nature 631, 449–458 (2024).

86. J. Dinnel, B. L. Moore, B. M. Skiver, P. Bose, Rasburicase in the management of tumor lysis: an evidence-based review of its place in therapy. Core Evid. 10, 23–38 (2015).

87. J. S. Sundy, H. S. B. Baraf, R. A. Yood, N. L. Edwards, S. R. Gutierrez-Urena, E. L. Treadwell, J. Vázquez-Mellado, W. B. White, P. E. Lipsky, Z. Horowitz, W. Huang, A. N. Maroli, R. W. Waltrip 2nd, S. A. Hamburger, M. A. Becker, Efficacy and tolerability of pegloticase for the treatment of chronic gout in patients refractory to conventional treatment: two randomized controlled trials: Two randomized controlled trials. JAMA 306, 711–720 (2011).

88. L. Du, Y. Zong, H. Li, Q. Wang, L. Xie, B. Yang, Y. Pang, C. Zhang, Z. Zhong, J. Gao, Hyperuricemia and its related diseases: mechanisms and advances in therapy. Signal Transduct. Target. Ther. 9, 212 (2024).

89. A. C. Nyborg, C. Ward, A. Zacco, B. Chacko, L. Grinberg, J. C. Geoghegan, R. Bean, M. Wendeler, F. Bartnik, E. O’Connor, F. Gruia, V. Iyer, H. Feng, V. Roy, M. Berge, J. N. Miner, D. M. Wilson, D. Zhou, S. Nicholson, C. Wilker, C. Y. Wu, S. Wilson, L. Jermutus, H. Wu, D. A. Owen, J. Osbourn, S. Coats, M. Baca, A therapeutic uricase with reduced immunogenicity risk and improved development properties. PLoS One 11, e0167935 (2016).

90. E. Oksanen, M. P. Blakeley, M. El-Hajji, U. Ryde, M. Budayova-Spano, The neutron structure of urate oxidase resolves a long-standing mechanistic conundrum and reveals unexpected changes in protonation. PLoS One 9, e86651 (2014).

91. E. Girard, S. Marchal, J. Perez, S. Finet, R. Kahn, R. Fourme, G. Marassio, A.-C. Dhaussy, T. Prangé, M. Giffard, F. Dulin, F. Bonneté, R. Lange, J. H. Abraini, M. Mezouar, N. Colloc’h, Structure-function perturbation and dissociation of tetrameric urate oxidase by high hydrostatic pressure. Biophys. J. 98, 2365–2373 (2010).

92. T. A. Hopf, A. G. Green, B. Schubert, S. Mersmann, C. P. I. Schärfe, J. B. Ingraham, A. Toth-Petroczy, K. Brock, A. J. Riesselman, P. Palmedo, C. Kang, R. Sheridan, E. J. Draizen, C. Dallago, C. Sander, D. S. Marks, The EVcouplings Python framework for coevolutionary sequence analysis. Bioinformatics 35, 1582–1584 (2019).

93. S.-H. Huang, T.-K. Wu, Modified colorimetric assay for uricase activity and a screen for mutant Bacillus subtilis uricase genes following StEP mutagenesis. Eur. J. Biochem. 271, 517–523 (2004).

## References and Notes

1. A. S. Parker, Y. Choi, K. E. Griswold, C. Bailey-Kellogg, Structure-Guided Deimmunization of Therapeutic Proteins. Journal of Computational Biology 20 (2), 152–165 (2013), doi:10.1089/cmb.2012.0251, 10.1089/cmb.2012.0251.

2. R. S. Salvat, Y. Choi, A. Bishop, C. Bailey-Kellogg, K. E. Griswold, Protein deimmunization via structure-based design enables efficient epitope deletion at high mutational loads. Biotechnology and Bioengineering 112 (7), 1306–1318 (2015), doi:10.1002/bit.25554, 10.1002/bit.25554.

3. X. Wang, et al., Diffusion language models are versatile protein learners. arXiv preprint arXiv:2402.18567 (2024).

